# Noncanonical Circular RNAs and Potential Functions

**DOI:** 10.64898/2026.07.23.740256

**Authors:** Kang Li, Weixu Wang, Abebe Edao Negesso, Jing Deng, Haoning Qin, Jiahao Jiang, Ke Ma, Jie Zhang, Ping Wei, Dawei Li, Feng-Ming (Spring) Kong, William C. Cho, Shulan Qiu, Weixiong Zhang

## Abstract

Circular RNAs (circRNAs) are ubiquitous in eukaryotes; dysregulated circRNA expression is linked to diseases, including lung cancer. In contrast to canonical circRNAs arising from exon-intron boundaries, noncanonical circRNAs originating within exonic, intronic, and intergenic regions have typically been dismissed as transcriptional noise or technical artifacts. To explore circRNA diversity and appreciate their functions, we developed an algorithm to identify both canonical and noncanonical circRNAs without relying on genome annotation, enabling the identification of circRNAs of all types and in newly sequenced or poorly annotated species. Results from lung cancer cells revealed that noncanonical circRNAs constituted over two-thirds of the circRNA population and were expressed more abundantly than canonical circRNAs, and genes with fewer and shorter exons were hotspots for noncanonical circRNA and circRNA isoform production. Further analyses showed that many noncanonical circRNAs were indeed endogenous circRNAs transcribed within cells rather than experimental artifacts, were potentially translated into proteins or peptides, and were conserved across species. Moreover, we validated 65 noncanonical circRNAs in NCI-H23 cells using multiple bioassays and demonstrated that both exonic and intergenic noncanonical circRNAs influenced cell viability. CircRNA profiles in tumor and tumor-adjacent tissues of lung cancer patients revealed tissue-specific expression and differentially expressed canonical and noncanonical circRNAs from cognate genes involved in cancer-related pathways, indicating their potential clinical relevance. This study confirmed the authenticity of noncanonical circRNAs and provided the first experimental evidence that noncanonical circRNAs influence cancer cell phenotypes. These findings broaden our understanding of circRNA biology, highlighting their widespread genomic distribution, diverse functions, and potential clinical relevance.

## 1. INTRODUCTION

Circular RNAs (circRNAs)^[1–3]^ constitute a distinct RNA species characterized by their single-stranded, covalently closed structure, which grants them greater stability than linear RNAs. Although circRNAs were first discovered decades ago^[4]^ and have been found to exist across nearly all eukaryotic lineages^[5]^, only in recent years have their biogenesis, functional roles, and clinical relevance started to be investigated and appreciated^[3]^. These circular molecules are now known to perform a variety of molecular functions, such as modulating transcription by interacting with transcription factors^[6]^, influencing mRNA and protein expressions through competition in RNA splicing^[7]^ and acting as competing endogenous RNAs (ceRNAs), also known as microRNA sponges, to regulate gene functioning in signaling pathways^[1, 2]^. Additionally, some circRNAs can be translated into polypeptides, further expanding their functional repertoire^[8]^. Notably, dysregulated circRNA expression has been associated with various diseases, including neurodegenerative disorders^[9]^, cardiovascular diseases^[10]^, and cancers^[11, 12]^. In lung cancer in particular, circRNA circPVT1 has been identified to promote cell proliferation by acting as a sponge for multiple microRNAs, including let-7, thereby modulating the expression of many cancer-related genes, such as KRAS^[13]^, which suggests circPVT1’s potential as a biomarker for lung cancer^[14]^.

Most reported circRNAs originate from genic regions of the genome, which consist of exons, forming exonic circRNAs (EcircRNAs)^[15]^, or a combination of exons and introns through intron retention, resulting in exonic-intronic circRNAs (EIcircRNAs)^[16]^. These circRNAs are back-splicing products of RNA splicing mechanisms^[17, 18]^ and utilize splicing donor and acceptor signals. Accordingly, the resulting circRNAs have their back-splicing junctions (BSJs) precisely aligned with exon-intron boundaries. Building on this model, over a dozen computational methods have been developed to predict circRNAs from RNA-seq data^[19]^. To distinguish true back-splicing events from sequencing artifacts, these algorithms typically rely on one of two rigid filtering strategies: they either require candidate junctions to strictly align with established exon-intron boundaries based on genome annotations, or they demand the presence of canonical splicing signals (i.e., AG/GT motifs) flanking the junctions. By discarding candidate junctions that fail to meet these specific structural or sequence prerequisites, these annotation- or signal-dependent approaches introduce a substantial algorithmic bias, primarily identifying *canonical* or *noncanonical circRNAs* derived from genic regions of the genome.

Empirical evidence has demonstrated that RNA circularization can occur at a broader range of genomic loci, with their BSJs occurring beyond exon-intron boundaries^[20]^. For instance, the oncogenic fusion circular RNA F-Circ2 is formed by fusing PML and RARα at loci that do not align with standard exon-intron boundaries^[21]^. Such *noncanonical* circRNAs can originate from genic regions, noncoding transcripts^[22, 23]^, and intergenic regions^[20, 24, 25]^. Here, we categorize these circRNAs into the class of *noncanonical circRNAs*, previously called interior circRNAs^[20]^. A significant number of noncanonical circRNAs have been identified in the genomes of two animal species (humans and mice) and one plant species (rice), and some have been experimentally validated in HeLa cells^[20]^. Additionally, noncanonical circRNAs have also been detected in normal and psoriatic skin^[26]^, further highlighting their presence and potential biological relevance.

Because they lack canonical splicing signals or fail to align with established genome annotation, noncanonical circRNAs are typically discarded as sequencing artifacts or transcription noise. Consequently, they are systematically excluded by conventional, genome-annotation-dependent identification pipelines^[27–29]^. In other words, no existing method can systematically identify all types of circRNAs and characterize their genomic origins, distributions, and sequence features. As a result, the full landscape of the circular transcriptome—encompassing both canonical and noncanonical types—remains incompletely mapped, leaving their potential impact on physiological processes, pathological conditions and diseases largely unexplored.

To address these limitations, we developed CAT, a *de novo* computational tool that comprehensively identifies both canonical and noncanonical circRNAs. Its core innovation lies in a genome-annotation-independent filtering strategy. Rather than using predefined exon-intron boundaries or canonical AG/GT splicing signals as prerequisites, CAT implements an objective filter driven solely by RNA-seq reads at candidate BSJs. This allows CAT to identify noncanonical circRNAs that conventional methods missed. To benchmark CAT’s performance, we first compared it with 16 existing pipelines for identifying canonical circRNAs^[19]^. We then applied CAT to comprehensively identify canonical and noncanonical circRNAs in lung cancer cells, examine their genomic distributions, classify them into subtypes, characterize their sequence features, and profile their expression abundance. By combining three orthogonal bioassays, PCR followed by Sanger sequencing for accessing BSJ sequences, RNase R treatment for removing linear RNAs, and targeted Amplicon sequencing (AmpliSeq), we experimentally validated 65 noncanonical circRNAs in lung cancer cells. To explore their cellular function, we investigated two noncanonical circRNAs in lung cancer cells using siRNA-mediated knockdown and the NanoString gene function analysis technique. Finally, we assessed the clinical relevance of noncanonical (and canonical) circRNAs in tumor and tumor-adjacent tissues from a cohort of lung cancer patients. Collectively, our findings demonstrate that noncanonical circRNAs are prevalent in cancer cells, constitutive members of the circRNA family, and involved in various cellular processes in lung cancer.

## 2. RESULTS

### 2.1. Circular RNA of All Types and the CAT Method

The BSJs of circRNAs from genic regions may have three possible positions relative to the nearest exon-intron boundaries of their cognate linear transcripts, giving rise to three circRNA subtypes (Figure 1A upper panel): (1) *canonical circRNAs* with both BSJs aligning to annotated exon-intron boundaries; (2) *half-boundary circRNAs* with one, but not both, of the BSJs aligning to exon-intron boundaries; and (3) *complete*- *nonboundary circRNAs* with both junctions appearing inside of exons or introns. *Lariats* are byproducts of splicing and can be regarded as variants of canonical circRNAs; for clarity, we list lariats separately here. *Intergenic circRNAs*, derived from intergenic regions that often encode poorly annotated noncoding transcripts, can be considered a new distinct subtype (Figure 1A upper panel). We collectively refer to complete-nonboundary, half-boundary, and intergenic circRNAs as *noncanonical circRNAs* (*nc-circRNAs*) to distinguish them from *canonical circRNAs* (*c-circRNAs*). By leveraging the back-splicing mechanism, c- circRNAs can be screened with relatively high accuracy using genome annotation, as most existing computational methods do^[1, 27–40]^. In sharp contrast, nc-circRNAs have not yet been fully detected, well profiled or characterized, even though many intronic half-boundary circRNAs have been reported^[22]^.

**Figure 1.**
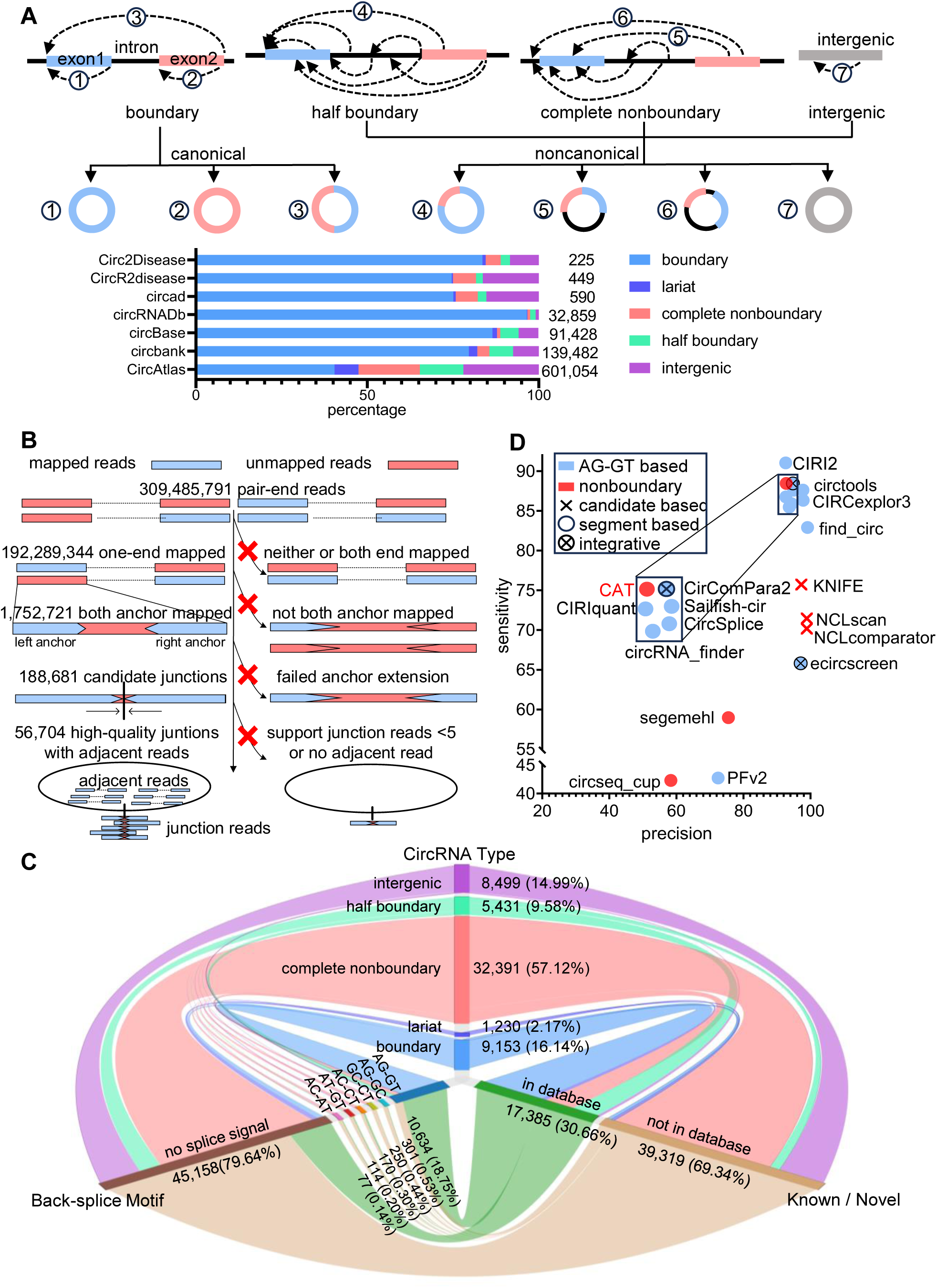
circRNA subtypes and their distributions and characteristics, and the CAT method and its performance. (A) ***circRNA subtypes***. Upper panel: A schematic view of different types of circRNAs and their genomic origins, including exons (orange/blue), introns (black), and intergenic regions (gray), where dashed curves indicate back-splicing events. Lower panel: The proportions of circRNA subtypes documented across seven public databases. (B) ***CAT workflow***. Left panel: Sequential RNA-seq read processing showing filtering statistics from the NCI- H23 dataset, starting with 309,485,791 input reads and resulting in 56,704 high-confidence circRNA candidates. Right panel: Schematic representation of mapping criteria at each step, including split-read mapping strategy, junction identification, and adjacent read support assessment. (C) ***Characteristics and relationships of circRNA subtypes*** *in NCI-H23 cells*. The relationships among subtypes (top), splice signals at BSJs (bottom-left), and presence in databases (bottom-right). (D) ***Performance benchmarking***. Comparison of CAT and 16 existing methods for finding c-circRNAs, in precision (x-axis) and sensitivity (y-axis). The methods are categorized by their key algorithmic features (indexed by graphic symbols). The inset box shows a magnified, better view of 6 methods with compatible performance.

To corroborate the authenticity of nc-circRNA subtypes, we analyzed them in existing circRNA repositories. Among 13 popular databases^[41–53]^, we selected 7 that provided unambiguous strand information for precise classification. Re-annotating the reported circRNAs in these databases revealed that all subtypes have already been documented (Figure 1A, lower panel). This meta-analysis of independent public data provided compelling evidence that nc-circRNAs are genuine circular transcripts rather than methodological artifacts, reinforcing the urgent need for their systematic investigation.

We followed and extended the circRNA nomenclature^[54]^ to name all types of circRNA. Canonical circRNAs were designated with the prefix “c-circ”followed by the host gene name, e.g., c-circCDYL for a c-circRNA from *CDYL*. Noncanonical circRNAs are designated with the prefix “nc-circ”, e.g., nc-circDNAJB6P1 for an nc- circRNA from *DNAJB6P1*. For multiple circRNAs from the same gene, numerical suffixes were added to their names based on the chromosomal loci of their 5’ junctions, e.g., nc-circPVT1-1 and nc-circPVT1-2. Intergenic nc-circRNAs, lacking associations with host genes, were named using the prefix “nc-circINT”followed by the chromosome number and a numerical identifier, e.g., nc-circINT2-1 and nc-circINT2-2 for two intergenic nc- circRNAs on chromosome 2.

To explore the overall landscape of circRNAs in the genome of any species, including those with newly sequenced or poorly annotated genomes, we developed a *de novo* approach called CAT (<u>C</u>ircRNA of <u>A</u>ll <u>T</u>ypes) to comprehensively and efficiently identify both canonical and noncanonical circRNAs (Figure 1B; see Methods and Supplemental Methods). The CAT pipeline supports both single-end and paired-end short reads, as well as single-cell RNA-seq data to identify and profile all types of circRNA at single-cell resolution.

The CAT method follows a three-step workflow (Figure 1B). First, sequencing reads are mapped to the reference genome, and unmapped reads are retained for circRNA identification. Second, every unmapped read is split-mapped to the genome; two short sequences (referred to as *anchors*, 30 bp long in our implementation) are extracted from the two read ends and independently mapped to the genome. The two genome-mapped anchors are then extended toward each other within the sequencing read and along the genome to detect the BSJs of a candidate circRNA (Figure 1B). Third, candidate BSJs are taken if they have supporting sequencing reads adjacent to the junctions.

#### 2.1.1. Canonical circular RNAs identified by CAT

We first evaluated the CAT method in identifying canonical or c-circRNAs. CAT’s performance on c-circRNAs can be used as a proxy for its performance on noncanonical or nc-circRNAs. For this analysis, we leveraged data and results from a recent benchmarking study^[19]^ that compared 16 existing circRNA detection tools^[1, 19, 27–40]^ (Supplemental Table S1). This comparative study generated a large circRNA-enriched, RNase R- treated RNA-seq dataset from NCI-H23 lung adenocarcinoma cells, providing over 309 million reads. It also experimentally validated 234 circRNAs in NCI-H23 cells using three orthogonal methods: PCR, RNase R treatment to enrich circRNAs followed by PCR, and Amplicon sequencing (Ampli-seq) for high-throughput targeted validation. Additionally, it compiled a list of circRNAs reported by the 16 benchmarked methods, which we used in our evaluation.

From the large NCI-H23 dataset, CAT identified 188,681 BSJs (Figure 1B). After applying CAT’s filtering criteria (i.e., minimal number of reads mapping to BSJs and enough reads adjacent to the BSJs; see Methods), 56,704 high-quality BSJs and their corresponding circRNAs were identified (Figure 1B), where 39,319 (69.34%) have not been documented in any of 13 public circRNA databases^[41–53]^ (Figure 1C). It is important to note that genome annotation was not used to identify these circRNAs, but rather helped classify them into subtypes, with the canonical subtype, c-circRNAs, accounting for 16.14% (9,153/56,704) of the total (Figure 1C;). Consistent with their origin from the canonical splicing machinery, over 87% of these c-circRNAs exhibited canonical splicing donor and acceptor signals at their junctions.

CAT’s highly reliable c-circRNA identification was further corroborated by existing public repositories. Specifically, 92.72% (8,487/9,153, Supplemental Figure S1A) of the detected c-circRNAs appeared in at least one circRNA database^[41–53]^, establishing a precision rate for CAT of no less than 92.72% (Figure 1D; Table S1). We then assessed sensitivity using the 234 experimentally validated circRNAs from the benchmarking study^[19]^ as a gold standard. CAT successfully detected 207 of these transcripts, demonstrating a sensitivity of at least 88.46% (Figure 1D; Table S1). When compared to the 16 existing methods, CAT achieved a robust balance; for instance, find_circ achieved the highest precision (99.13%) but lower sensitivity (82.91%), whereas CIRI2 reached the highest sensitivity (91.03%) with slightly lower precision (92.60%). It is crucial to note that this comparison inherently favored the established methods, as the experimentally validated circRNAs were originally selected from their prediction pools, excluding any unique CAT predictions. Despite this bias, CAT identified 664 novel c-circRNAs (7.25% of the 9,153 c-circRNAs) that were completely missed by all 16 existing pipelines (Table S2).

In short, CAT excels as a top-performing method for detecting c-circRNAs, achieving high sensitivity and precision (Figure 1D, Table S1). While CIRI2 and CirComPara2 are good alternatives for c-circRNAs, they are unable to find nc-circRNAs. Furthermore, as CAT operates independently of genome annotation, it can be applied to species with either no or poorly annotated genomes. Because CAT adopts the same strategy for both c-circRNAs and nc-circRNAs, its performance on c-circRNAs serves as a reliable proxy for its capability in identifying nc-circRNAs.

#### 2.1.2. Noncanonical circular RNAs identified by CAT

CAT identified 46,321 nc-circRNAs, 81.69% of all 56,704 circRNAs identified from the NCI-H23 dataset under a read cutoff of 5. These nc-circRNAs included 32,391 (57.12% of all circRNAs) complete nonboundary, 5,431 (9.58%) half boundary, and 8,499 (14.99%) intergenic circRNAs (Figure 1C). In sharp contrast to the c- circRNAs, these noncanonical subtypes exhibited fundamentally different sequence signatures. Fewer than 6% of complete nonboundary and intergenic circRNAs, and less than 25% of half boundary circRNAs, carried known splicing signals next to their BSJs (Figure 1C; Supplement Figure S1B).

Interestingly, although the existing methods are not designed for nc-circRNAs, 4,055 (12.52% of 32,391) complete nonboundary, 3,726 (68.61%) half-boundary, and 448 (5.27%) intergenic circRNAs have been documented across 13 databases (Figure 1C), supporting the authenticity of nc-circRNAs. The vast majority (87.48% of complete nonboundary and 94.73% of intergenic circRNAs) are novel and have not been previously reported. This stark contrast indicates that existing methods have largely overlooked this dominant population of circular transcripts.

CAT can also effectively identify all types of circRNA in single-cell or single-nucleus RNA-seq (scRNA-seq or snRNA-seq) data, provided that their cDNA libraries are constructed with circRNAs retained, like using random primers but not Poly-A library construction protocol that excludes circRNAs. As a demonstration, we applied CAT to a mixed total RNA sample comprising lung cancer cells (A549 and HCC827) and leukemia cells (K562), which were sequenced using barcoded semi-random primers to differentiate cells. After preprocessing for quality control and cell sorting, CAT identified both canonical and noncanonical circRNAs of different cell types (Figure S1C, S1D; see Methods).

### 2.2. Characteristics of Noncanonical (and Canonical) Circular RNAs

Deep profiling of circRNAs in NCI-H23 cells revealed that nc-circRNAs outnumbered c-circRNAs by a 4:1 ratio (Figure 1C; where lariats and c-circRNAs count for 18.31%), and more nc-circRNAs were distributed across all 23 chromosomes compared to c-circRNAs (Figure 2A). Interestingly, there appeared to be no correlation between the number of circRNAs and chromosome length; for example, chromosomes 1 and 19 carry comparable numbers of nc-circRNAs.

**Figure 2.**
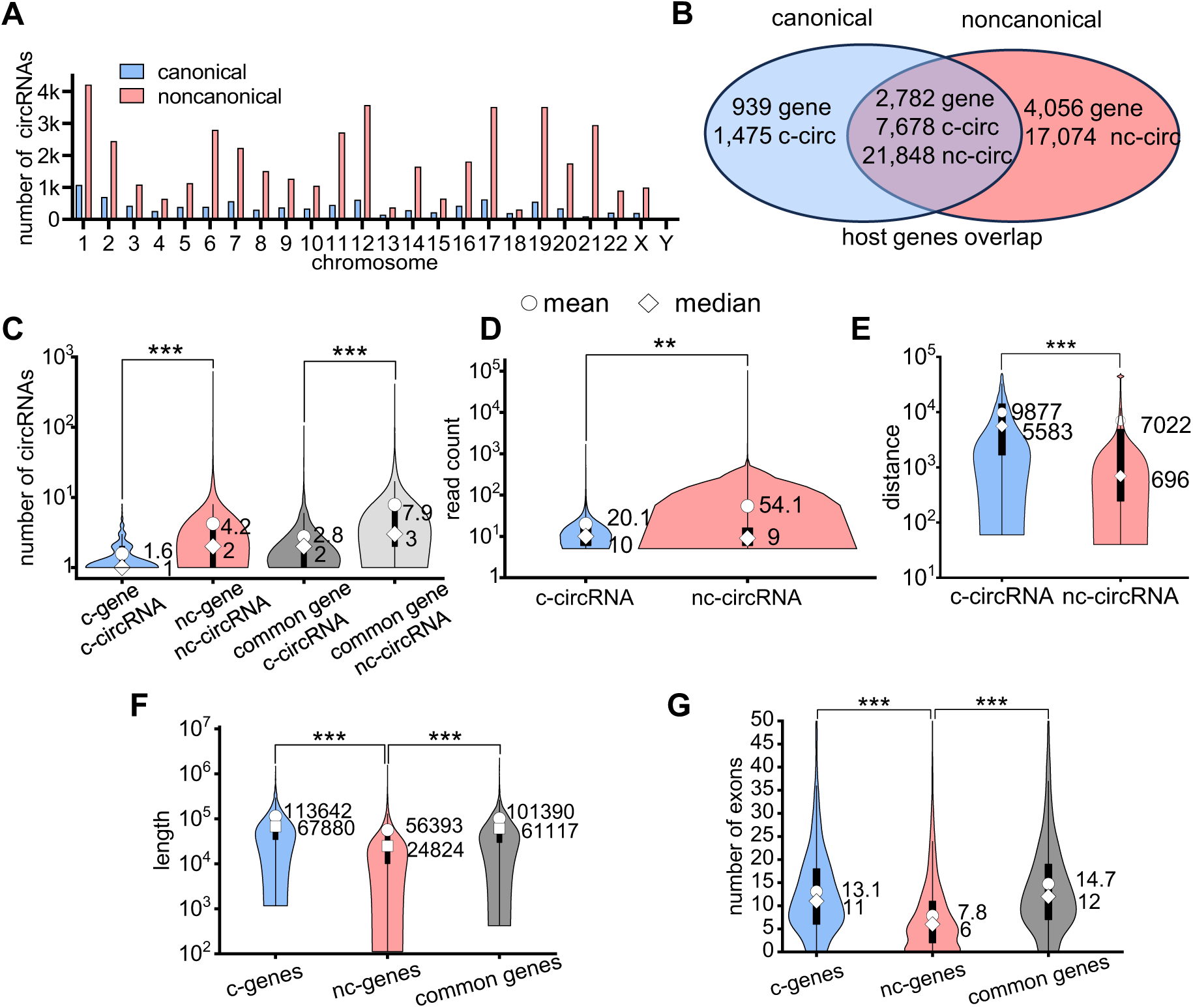
Genomic and expression features of nc-circRNAs versus c-circRNAs. (A) Distributions of c-circRNAs (light blue) and nc-circRNAs (pink) across the human chromosomes, showing no correlation between the numbers of circRNAs and the chromosome lengths. (B) nc-circRNAs and c-circRNAs from mRNA genes – there exist genes that generate exclusively nc-circRNAs (light blue, nc-circRNA-genes or nc-genes for shorts), exclusively c-circRNAs (pink, c-circRNA-genes or c- genes), and both (grey, common-genes). (C) nc-circRNAs are more numerous than c-circRNAs in the three types of host genes listed in (B), analyzed using an unpaired two-tailed t-test. (D) nc-circRNAs are expressed more abundantly than c-circRNAs, with expression abundance measured by the number of reads covering their BSJs (in log scale). (E) Distributions of the distances of the two loci of circRNA BSJs (in log scale), indicating nc-circRNAs are typically shorter than c-circRNAs. (F) mRNA genes hosting exclusively nc-circRNAs are statistically shorter and (G) they have fewer exons than the other two host gene types listed in (B).

A striking feature of circRNAs was that multiple circRNAs could arise from a single host transcript, constituting circRNA isoforms. We examined this feature on the complete-nonboundary and half-boundary nc-circRNAs from genic regions, which encompass nearly 70% of nc-circRNAs in NCI-H23. Specifically, 6,838 and 3,721 genes produced 38,922 nc-circRNAs and 9,153 c-circRNAs, respectively, indicating that circRNA isoforms arise from the same loci. Among all 7,777 host genes, 35.77% (2,782/7,777) generated both c- circRNAs and nc-circRNAs, which we referred to as *common genes* for convenience (Figure 2B, overlapping area). These common genes accounted for 83.89% of c-circRNA and 56.13% of nc-circRNA productions, suggesting a tendency for genes producing c-circRNAs to also generate nc-circRNAs. In contrast, 939 genes produced exclusively 1,475 c-circRNAs (*c-circRNA genes*), whereas 4,056 genes generated exclusively 17,074 nc-circRNAs (*nc-circRNA genes*). Common genes hosted more circRNAs, with c-circRNA-to-gene and nc- circRNA-to-gene ratios of 2.76 and 7.85, respectively, surpassing those of c-circRNA genes (1.57) and nc- circRNA genes (4.21).

Remarkably, nc-circRNAs from nc-circRNA genes were not only more numerous per gene (Figure 2B, 2C) but also exhibited higher average expression abundance than c-circRNAs from c-circRNA genes (Figure 2D), as determined by the number of reads spanning their BSJs. Notably, nc-circRNA genes and their nc-circRNAs were statistically shorter than c-circRNA genes and their c-circRNAs (Figure 2E, 2F). Additionally, nc-circRNA genes contained significantly fewer exons compared to c-circRNA genes (Figure 2G).

Together, these findings suggest that short nc-circRNA genes constitute genomic hotspots for nc-circRNA production, and that nc-circRNAs are more abundant than c-circRNAs in both numbers and expression levels.

The possible association of circRNA subtypes with specific spliceosomes, particularly the major (U2- dependent) and minor (U12-dependent) spliceosomes, is an important feature to study; the results hint at their biogenesis. Utilizing the Intron Annotation and Orthology Database (IAOD)^[55]^, we grouped all circRNAs detected in NCI-H23 cells^[19]^ into four categories (see Methods): containing exclusively U2 signals, exclusively U12 signals, both signals, or neither (Supplementary Table S3). As anticipated, c-circRNAs relied almost entirely on the major spliceosome, with 98.57% (9,022/9,153) carrying exclusively U2 signals and an additional 0.07% (6/9,153) harboring both U2 and U12 signals. Half-boundary circRNAs and lariats, subtypes carrying one exon-intron boundary, exhibited high U2-signal occurrence of 79.30% (4,307/5,431) and 72.85% (896/1,230), respectively. Interestingly, however, no complete-nonboundary circRNAs carried U2 signals, and only 0.02% (8/32,391) of them were associated with U12 signals; likewise, no intergenic circRNAs had any U2 or U12 signatures. These results suggest that the two subtypes may originate from distinct biogenesis mechanisms, a subject for future research.

### 2.3. Noncanonical CircRNAs are Authentic Circular RNAs

Several experiments were conducted to provide multiple lines of evidence supporting that nc-circRNAs are authentic circRNAs. The results from these bioassays and genomic analysis also helped reveal their cellular functions.

#### 2.3.1. Validation by multiple bioassays

We experimentally validated CAT’s identified nc-circRNAs in NCI-H23 cells, focusing on confirming their BSJ authenticity, a hallmark of circRNAs, using Amplicon sequencing (Ampli-seq), PCR, PCR followed by Sanger sequencing, and RNase R treatment followed by qPCR. We designed divergent primers spanning BSJs and convergent primers as negative controls for linear RNAs (Figure 3A). PCR with cDNA and gDNA templates ensured that circRNAs were RNA-specific and not derived from genomic DNA or contamination. Ampli-seq enabled high-throughput, targeted BSJ detection, and Sanger sequencing provided high-confidence BSJ sequences and often full-length circRNA sequences directly. RNase R digestion degraded linear RNAs, enriching circular RNAs, followed by qPCR to confirm circRNA presence post-treatment.

**Figure 3.**
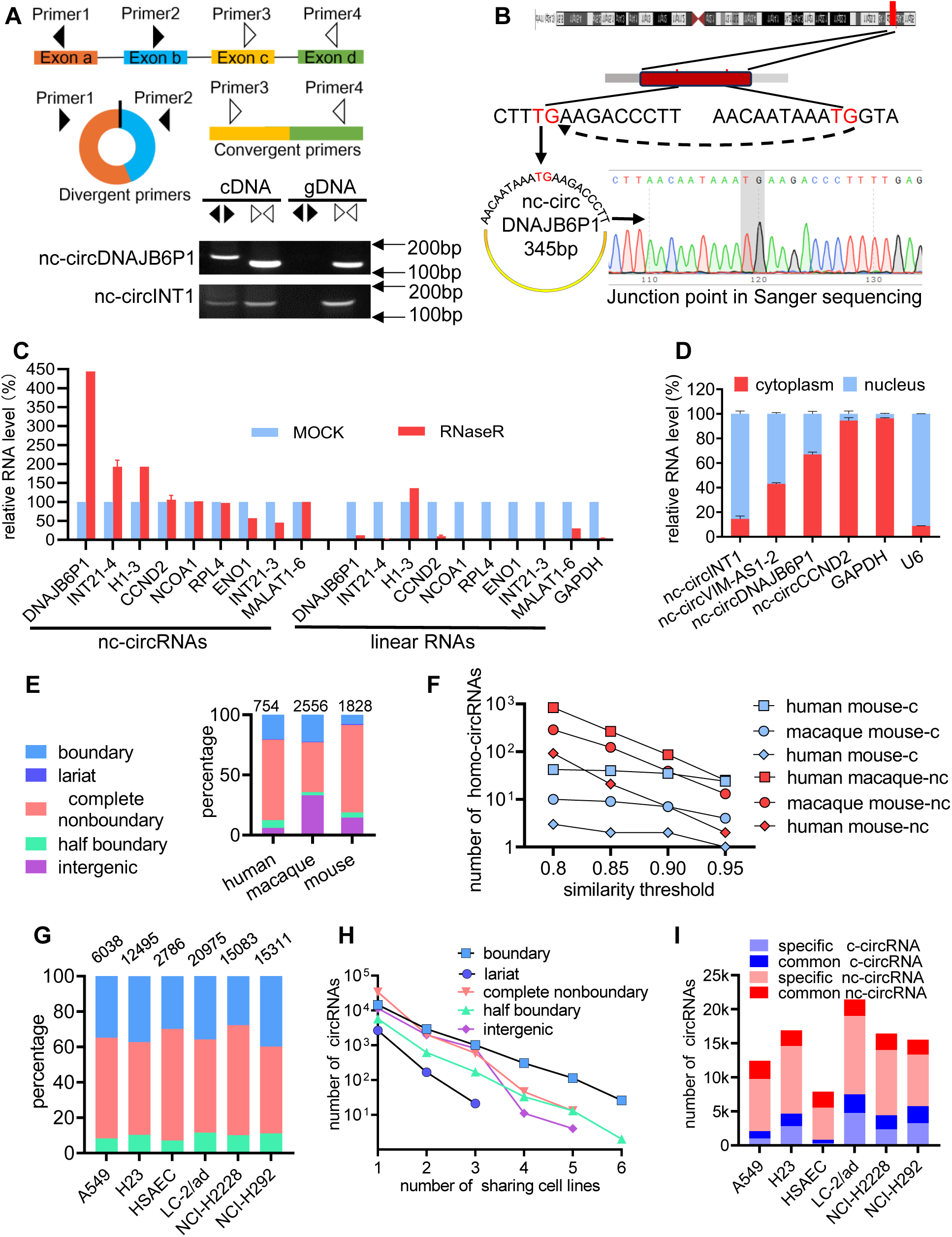
Experimental validation and conservation analysis of nc-circRNAs. (A) ***PCR-based validation***. Upper: Primer design scheme where divergent primers for circRNA detection and convergent primers for linear RNA detection as a control. Lower: PCR validation of three representative nc- circRNAs using divergent and convergent primers with cDNA and gDNA templates. (B) ***Sanger sequencing*** of nc-circDNAJB6P1. Shown from the top are genomic location and structure, BSJ sequences and a sequencing chromatogram confirming the BSJ. (C) ***RNase R treatment assays*** followed by qPCR on some nc-circRNAs and their linear counterparts in RNase R-treated (red) and mock-treated (light blue) samples. Data normalized to mock treatment, shown as mean ± SEM. Statistical significance is indexed by asterisks: * p < 0.05, ** p < 0.01, *** p < 0.001. (D) ***Subcellular localization analysis***. qPCR quantification of 4 nc-circRNAs in cytoplasm (red) and nucleus (blue), with GAPDH and U6 as compartment-specific controls. Data are shown in percentage of total RNA. (E, F) ***Sequence conservation*** of circRNA subtypes across humans, macaques, and mice (using lung tissues). (E) ***Relative proportions*** of different circRNA categories in the lung tissues of the three species. (F) ***Conservation*** of circRNAs between species pairs. Shown are the numbers of conserved circRNAs between two species at different sequence similarity thresholds (0.8-0.95) within a 300-nt window around BSJs. Suffices ‘nc’ and ‘c’ indicate nc-circRNA and c-circRNA, respectively. (G-I) ***Expression conservation*** of subtypes across five lung cancer cell lines and a normal lung epithelial line (HSAEC). (G) ***Relative distributions*** of subtypes in six cell lines, showing that complete-nonboundary and intergenic nc-circRNAs are the most abundant. (H) ***Expression conservation across cell lines***. The number of circRNAs (y-axis, log scale) versus the number of cell lines in which they are expressed (x-axis). c-circRNAs are the most conserved, whereas nc-circRNAs tend to be cell-line-specific. (I) ***Cell specificity***. Cell line-specific nc-circRNAs/c-circRNAs versus nc-circRNAs/c-circRNAs in more than one cell line, showing more cell-specific nc-circRNAs than c-circRNAs.

A total of 102 CAT-predicted circRNAs were selected for validation, comprising 82 nc-circRNAs (59 complete- noncanonical, 1 half-boundary, and 22 intergenic circRNAs), 10 c-circRNAs, and 10 lariats (Table S4). These circRNAs were chosen for their potential functional relevance to lung cancer and their representation of circRNA subtypes.

Ampli-seq detected 62 (75.61%) of the 82 nc-circRNAs, 9 (90.00%) of the 10 c-circRNAs, and 6 (60.00%) of the 10 lariats (Table S4), achieving an overall validation rate of 74.51%. This is comparable to the validation rate for c-circRNAs in the benchmarking study^[19]^. Next, we validated a subset of the 102 circRNAs, including many missed by Ampli-seq, using PCR, PCR followed by Sanger sequencing, and RNase R treatment followed by qPCR. PCR detected 14 of 62 tested, including nc-circDNAJB6P1, and nc-circINT1 (Figure 3A; Table S4). Agarose gel electrophoresis of PCR products (Figure 3A) confirmed that these circRNAs originated from RNA (cDNA) rather than DNA (gDNA). Divergent and convergent primers confirmed that the RNA was circular rather than linear (Figure 3A). Sanger sequencing was successful for 5 of 12 circRNA PCR products, including nc-circDNAJB6 (Figure 3B). RNase R digestion followed by qPCR on circular and linear RNA determined the presence of 17 of 42 circRNAs, including nc-circDNAJB6P1 (Figure 3C; Figure S1E; Table S4).

Overall, 82 (80.39%) of the 102 circRNAs and 65 (79.29%) of the 82 nc-circRNAs were validated by at least one bioassay (Table S4). These results confirmed that nc-circRNAs are indeed authentic circRNAs, and indicated that CAT’s specificity is at least 80.39% for identifying all types of circRNA.

Moreover, several nc-circRNAs (nc-circDNAJB6P1, nc-circINT1, nc-circCCND2, and nc-circVIM-AS1-2) in NCI- H23 cells could be localized to the nucleus, cytoplasm, or both, as detected by qPCR on RNA from separated cytoplasmic and nuclear fractions (Figure 3D). Intergenic nc-circINT1 was primarily nuclear, exonic nc- circCCND2 was mainly cytoplasmic, and nc-circDNAJB6P1 and nc-circVIM-AS1-2 were detected in both compartments (Figure 3D).

#### 2.3.2. Direct native RNA-seq confirms nc-circRNAs are endogenous transcripts inside the cell

The results above robustly authenticated nc-circRNAs. Nevertheless, an inherent technical issue is template-switching artifacts^[56, 57]^ introduced by reverse transcriptase (RT) or PCR amplification during RNA-seq library preparation, which could generate spurious chimeric transcripts and possibly nc-circRNAs. To examine this technical bias, we leveraged orthogonal dRNA-seq data from Oxford Nanopore direct RNA sequencing across HEK293, HeLa and HepG2 cells^[58]^, which processes native RNA molecules directly through the nanopore while displacing the complementary cDNA strands, thereby avoiding RT- or PCR-induced template-switching artifacts.

While the method in [58] was not designed for circRNAs, we extracted 1,273 candidate circRNAs, among which 175 (13.7%) were intergenic, 19 (1.5%) complete-nonboundary, 3 half-boundary (0.2%), and 1 lariat (0.1%) nc-circRNAs (Table S5). Detecting these noncanonical junctions in direct, RT-free single-molecule sequencing data provided additional direct evidence that nc-circRNAs were endogenous transcripts rather than spurious experimental artifacts.

#### 2.3.3. Potential to be translated into proteins or peptides

Further evidence supporting nc-circRNA authenticity came from their potential to be translated into proteins or peptides^[59, 60]^. To explore this property, we applied stringent criteria and leveraged Ribo-seq data from A549 cells^[61]^. First, to maximize the utility of the Ribo-seq data, we sequenced in-house RNase R-treated RNA from A549 cells and applied the CAT method to identify all types of circRNAs, yielding 11,314 nc- circRNAs and 1,701 c-circRNAs (Table S6). Next, we assessed whether these 13,015 circRNAs contained circular Open Reading Frames (c-ORFs), a key indicator of translatability^[62]^, revealing that 39.30% (4,446/11,314) of nc-circRNAs and 89.18% (1,517/1,701) of c-circRNAs contained ORFs. We then performed an Internal Ribosome Entry Site (IRES) analysis^[63]^, showing that 48.36% (5,471/11,314) of nc-circRNAs and 78.31% (1,332/1,701) of c-circRNAs contained IRES entry points, suggesting their capacity to initiate translation. Finally, we analyzed the circRNAs against the Ribo-seq data from A549 cells^[61]^ to identify those that were actively associated with Ribosomes. Under this stringent criterion, 0.45% (51/11,314) of nc- circRNAs and 0.06% (1/1,701) of c-circRNAs were detected to carry Ribosome-protected fragments. One explanation for these low detection rates is that the conditions used in these experiments may not be compatible.

The integration of these three independent analyses revealed intriguing patterns in the potential coding capacity of nc-circRNAs. Among all circRNAs, 3,565 (27.39%) possessed both ORFs and IRES elements, suggesting their structural readiness for translation. However, only 31 circRNAs (0.24%) showed both ORF presence and evidence of Ribosome engagement, while 25 circRNAs (0.19%) demonstrated both IRES features and Ribosome occupancy. Although only a small fraction of circRNAs may be potentially translated into proteins or peptides, as reported previously^[64]^, we identified 13 nc-circRNAs and 1 c-circRNA (Table S7) that met all three criteria–containing ORFs, possessing IRES entry points, and showing evidence of Ribosome engagement. This highly selected group, representing 0.11% of nc-circRNAs and 0.06% of c-circRNAs, constituted the highest-confidence set of potentially coding circRNAs, underscoring their potential functional significance.

#### 2.3.4. Conservation across species

Gene conservation across species is a strong indicator of functional importance, as it suggests that evolutionary pressure has led to the retention of these elements. Therefore, the conservation of nc-circRNAs supports their authenticity. To investigate this, we analyzed circRNA conservation (see Supplemental Methods) in lung tissues from humans, macaques, and mice using public RNA-seq data [45], which yielded 22,254,764; 14,217,189; and 17,876,804 reads, respectively. The result showed that complete-nonboundary circRNAs were the dominant subtype across all three species (Figure 3E), consistent with the observations in NCI-H23 and A549 lung cancer cells.

Given the evolutionary divergence among the three species, assessing the conservation of circRNA is challenging. To address this issue, we evaluated conservation across the range of 80% to 100% sequence similarity within a 50 bp window around the BSJs of circRNAs. Both noncanonical and canonical circRNAs showed substantial similarities in pairwise comparisons (Figure 3F). Notably, while humans and mice had comparable proportions of complete noncanonical circRNAs, these circRNAs were more conserved between humans and macaques (Figure 3F), reflecting their closer evolutionary relationship. Overall, these findings provide further evidence that nc-circRNAs are genuine and functionally significant members of the circRNA family.

#### 2.3.5. Conservation across lung carcinoma cells

Furthermore, we investigated the conservation of nc-circRNA expression across five lung cancer cell lines (A549, NCI-H23, LC-2/ad, H2228, and H292) and a normal epithelial cell line (HSAEC). To this end, we generated in-house RNase R-treated random-primer-captured RNA-seq data from A549, NCI-H23, and HSAEC cells, and incorporated exome-capture RNA-seq data for the remaining LC-2/ad, H2228, and H292 cells from a study of circRNA in cancer^[11]^.

As expected, exome-capture RNA-seq profiling excluded nearly all circular transcripts from intergenic regions, which were missing from the LC-2/ad, H2228, and H292 cells (Supplemental Table S8). In concordance with earlier results, complete-nonboundary circRNAs were more abundant than c-circRNAs; the distributions of circRNAs from genic regions, where protein-coding genes reside, were compatible in these six cell lines (Figure 3G).

The patterns of the conserved or shared circRNAs across the six cell lines revealed that c-circRNAs were the most conserved, followed by complete-nonboundary, half-boundary, and intergenic nc-circRNAs, with lariats showing the least conservation (Figure 3H). Therefore, many nc-circRNAs and c-circRNAs showed conserved expression across different lung cancer cell lines, confirming their authenticity and suggesting their potential contribution to disease pathology.

A closer examination of cell line-specific versus common circRNAs revealed that nc-circRNAs were more likely to be cell type-specific, whereas c-circRNAs were more prevalent across multiple cell lines (Figure 3I). This pattern suggests that nc-circRNAs could potentially serve as biomarkers for specific lung cancer cell types.

### 2.4. Potential Functions in Lung Cancer

We performed several analyses to explore nc-circRNAs’ functions

#### 2.4.1. CircRNA host genes are enriched in cancer pathways

To infer circRNAs’ potential functions in lung adenocarcinoma, we first analyzed their genomic origins and host gene functions in NCI-H23 cells. More than 85% of detected circRNAs originated from genic regions (Figure 1C). Using host genes as a proxy, Kyoto Encyclopedia of Genes and Genomes (KEGG) analysis revealed that host genes of both nc-circRNAs and c-circRNAs were strongly enriched in cancer-related pathways, including cell cycle and survival (Figure 4A). These results align with the malignant nature of NCI-H23 cells and suggest broad involvement of circRNAs in lung cancer biology.

**Figure 4.**
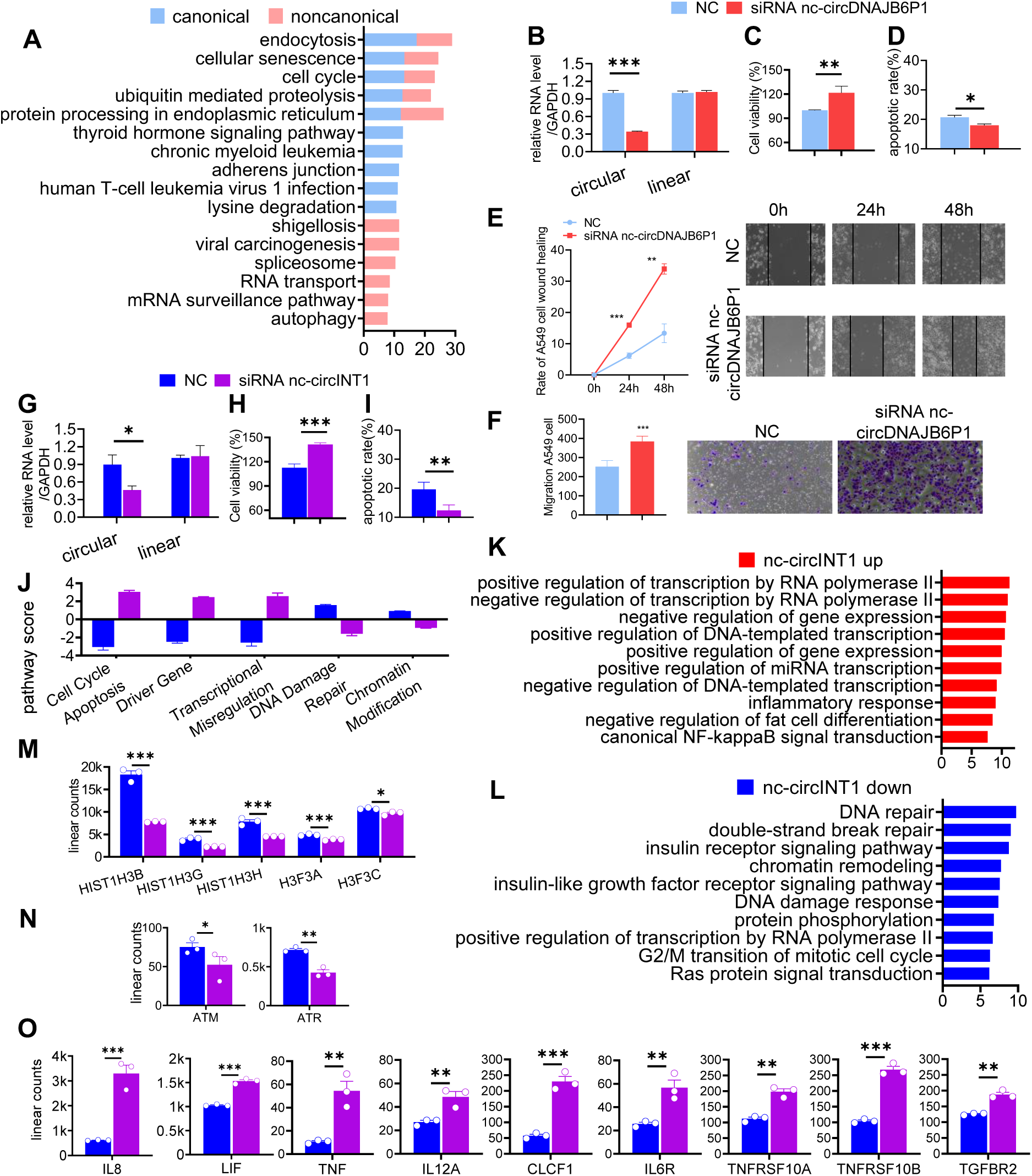
Cellular functions of nc-circRNAs in lung cancer cells. (A) ***Pathway enrichment*** of the host genes of nc-circRNAs and c-circRNAs in NCI-H23 cells. Bar length indicates statistical significance (-log10 *p*-value). (B-D) ***Functional analysis* of nc-circDNAJB6P1**. Data shown in mean ± SEM. Statistical significance indicated by asterisks: * p < 0.05, ** p < 0.01, *** p < 0.001. (B) ***qPCR*** on siRNA-mediated knockdown efficiency of the circular and linear forms. (C) ***Cell viability*** assays following knockdown. (D) ***Apoptosis analysis by flow cytometry*** after knockdown. (E, F) ***Knockdown assays*** of nc-circDNAJB6P1 enhances migration in A549 cells, as shown by wound healing (E) and Transwell assays (F). Both panels include representative images and quantification. (G-I) ***Functional analysis of nc-circINT1***. (G) ***qPCR*** on siRNA-mediated knockdown efficiency for circular and linear forms. (H) ***Cell viability*** assays following knockdown. (I) ***Apoptosis analysis by flow cytometry*** after knockdown. (J) ***Pathway analysis from NanoString*** data comparing effects of nc-circINT1 knockdown on major cellular pathways. Scores indicate pathway activation (positive) or suppression (negative). (K, L) ***Gene Ontology (GO) enrichment analysis*** of differentially expressed genes (DEGs) after nc-circINT1 knockdown. Statistical significance in log10(p-value) of DEG-affected biological processes is in x-axis. Red and blue bars represent pathways enriched with up- and down-regulated genes, respectively. (M) ***Downregulation of Histone H3*** genes; (N) ***DNA damage response genes***; and (O) ***Upregulation of inflammatory cytokine***s after nc-circINT1 knockdown.

#### 2.4.2. A pseudogene-derived nc-circRNA suppresses oncogenic phenotypes

Although host gene analysis suggested general cancer relevance, the specific functions of nc-circRNAs, especially from unconventional loci, remain poorly defined. We therefore investigated nc-circDNAJB6P1, an exonic circRNA derived from the pseudogene DNAJB6P1, a paralog of DNAJB6, a tumor suppressor implicated in breast cancer metastasis^[65]^. Pseudogenes have been reported to regulate cancer progression through diverse mechanisms, including miRNA sponging and antisense transcription^[65]^.

We confirmed the full-length structure and BSJ of nc-circDNAJB6P1 by PCR and Sanger sequencing (Supplementary Figure S1F). A BSJ-targeting siRNA specifically depleted the circular form (Figure 4B). Functional assays showed that nc-circDNAJB6P1 knockdown increased A549 cell viability (Figure 4C), reduced cell death (Figure 4D), and enhanced migration, as measured by wound healing and transwell assays (Figure 4E, F). Together, these data identified nc-circDNAJB6P1 as a pseudogene-derived circRNA that potently suppressed oncogenic phenotypes in lung cancer cells.

#### 2.4.3. An intergenic nc-circRNA regulates cell cycle, DNA damage, and inflammation

To further investigate nc-circRNAs from unconventional loci, we focused on nc-circINT1, a validated intergenic circRNA. Like nc-circDNAJB6P1, siRNA-mediated knockdown of nc-circINT1 (Figure 4G) promoted NCI-H23 cell viability and suppressed apoptosis (Figure 4H, I), supporting a tumor-suppressive role.

To define its molecular functions, we performed NanoString nCounter transcriptome profiling of NCI-H23 cells after nc-circINT1 knockdown. Principal component analysis showed clear separation between depleted and control cells (Supplementary Fig. S1H). Knockdown induced significant transcriptional changes (78 upregulated and 116 downregulated genes; Supplementary Fig. S1H, Table S9). Pathway score analysis indicated enhanced cell cycle – apoptosis and transcriptional misregulation pathways but suppressed DNA damage repair and chromatin modification (Figure 4J).

Gene Ontology (GO) analysis revealed that upregulated genes were enriched in transcriptional regulation and inflammatory responses (Figure 4K), whereas downregulated genes were involved in DNA damage response and repair (Figure 4L). Notably, replication-dependent histone H3.1 genes (HIST1H3B, HIST1H3G, HIST1H3H) and replication-independent histone H3.3 genes (H3F3A, H3F3C) were suppressed (Figure 4M), suggesting impaired chromatin dynamics and genomic instability. However, cell death was avoided (Figure 4H, I), likely due to downregulation of DNA damage sensors such as ATM and ATR (Figure 4N). In parallel, knockdown activated a pro-survival inflammatory program, including cytokines and chemokines (IL8, LIF, TNF, IL12A, CLCF1) and their receptors (IL6R, TNFRSF10A/B, TGFBR) (Figure 4O), along with NF-κB, JAK-STAT, MAPK, and PI3K signaling (Supplementary Fig. S2A). Conversely, Ras and Wnt pathways were suppressed (Supplementary Fig. S2A). The differentially expressed genes (DEGs) identified by NanoString, along with the corresponding GO terms, are listed in Supplementary Table S10.

Collectively, these findings establish nc-circINT1 as a regulator of chromatin integrity, DNA damage signaling, and inflammatory responses. This represents the first functional characterization of an intergenic circRNA in cancer.

In summary, this comprehensive analysis, which combines siRNA-mediated knockdown assays on individual nc-circRNAs with systematic profiling of their impacts on downstream genes, provides new insights into the molecular mechanisms underlying the functional roles of nc-circRNAs in cancer cells. Notably, we demonstrated for the first time that novel types of circRNAs, specifically exonic and intergenic nc-circRNAs such as nc-circDNAJB6P1 and nc-circINT1, can significantly influence cell viability, cell migration, chromatin and DNA integrity, as well as trigger inflammatory responses when knocked down. These findings highlight the previously underappreciated functional diversity of circRNAs, extending beyond their roles as miRNA sponges, and reveal their involvement in both fundamental cellular processes and potential pathological pathways, such as inflammation and DNA damage response in lung cancer. Our results emphasize the importance of further studies of nc-circRNAs as key regulators in cellular physiology and disease.

### 2.5. Potential Relevance to Lung Cancer

We investigated the potential clinical relevance of nc-circRNAs, along with c-circRNAs, in a small cohort of eight patients with non-small cell lung cancer (NSCLC)^[66]^. A differential expression of their RNase R-treated and random-primer-captured RNA-seq datasets of paired tumor and tumor-adjacent tissues identified 189,049 unique circRNAs. Classification of these circRNAs put them into 126,472 (66.9%) complete- nonboundary, 40,564 (21.5%) canonical, 14,072 (7.4%) half-boundary, 6,367 (3.4%) intergenic, and 1,574 (0.8%) lariat subtypes (Figure 5A, the first bar), compatible with the distribution in NCI-H23 cells (Figure 1C).

**Figure 5.**
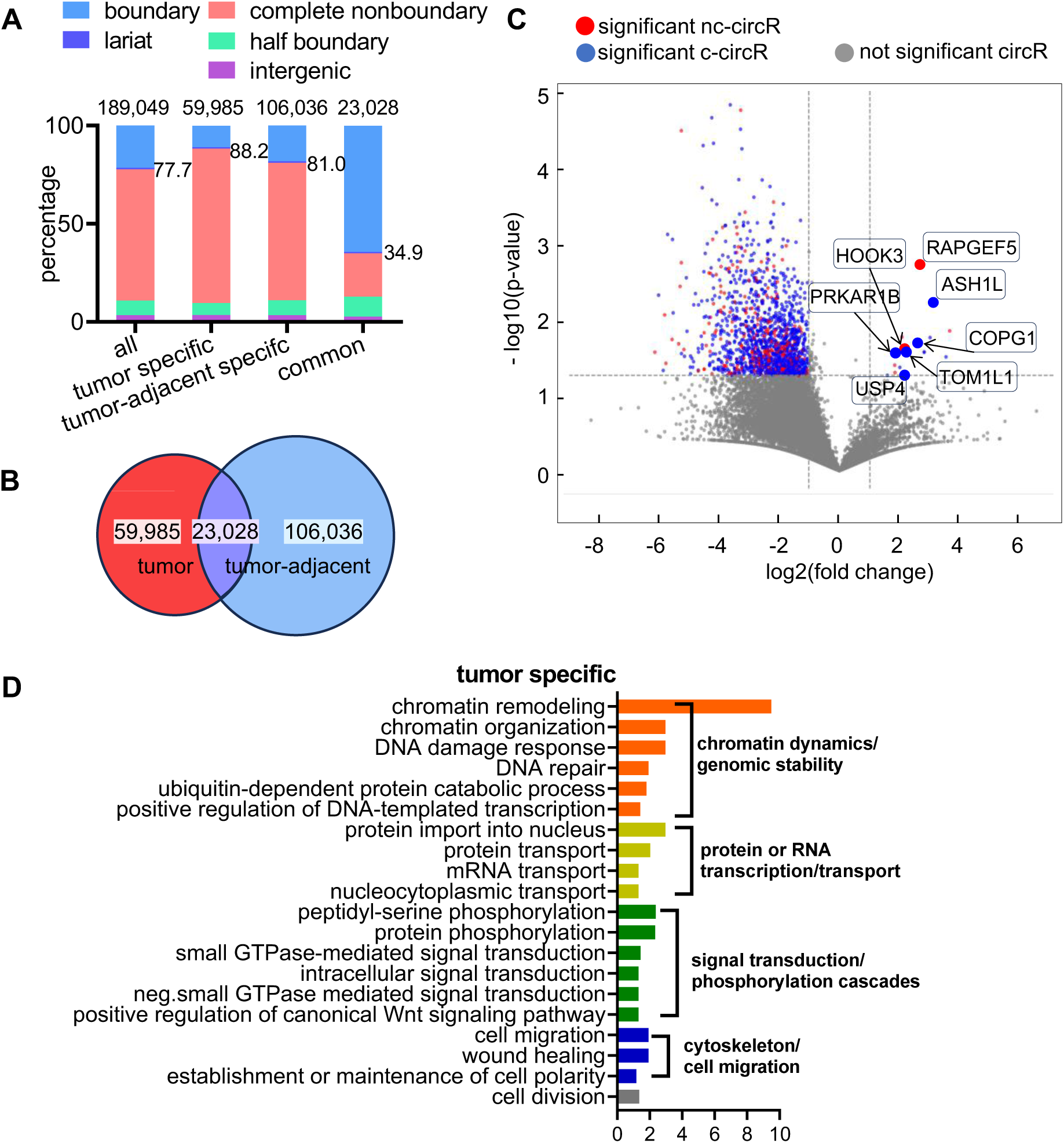
nc-circRNAs exhibit tissue-specific expression patterns and potential relevance in NSCLC. (A) Distributions of all subtypes of circRNAs (first bar), tumor-specific circRNAs (second bar), tumor-adjacent specific circRNAs (third bar), and circRNAs in both tissue types (fourth bar). The number of unique circRNAs is on the top of each bar and the number in the middle is the percentage of all nc-circRNAs. (B) Distribution of 189,049 unique circRNAs in tumors, tumor-adjacent tissues and both tissues of NSCLC. (C) Differential expressions of circRNAs expressed in both tumors and tumor-adjacent tissues. Representative upregulated circRNAs are annotated with their host gene names to highlight their known involvement in cancer. Dashed lines indicate significance thresholds (|log2(FC)| > 1, p < 0.05). (D) GO enrichment of host genes of 1,283 tumor-specific circRNAs. Enriched biological processes are grouped into four major functional categories listed on the right. The x-axis represents the −log10(p-value) of enriched terms.

These unique circRNAs were distributed markedly differently across the tumor and tumor-adjacent tissues. More than half, i.e., 56.1% (106,036), arose exclusively in tumor-adjacent samples; 31.7% (59,985) appeared only in tumors; and 12.2% (23,028) were shared between the two tissue types (Figure 5B), indicating tissue- specificity. Indeed, nc-circRNAs showed greater tissue specificity. They comprised 77.7% of the overall 189,049 unique circRNAs (Figure 5A, the first bar), increasing to 88.2% and 81.0% exclusively in tumors and in tumor-adjacent tissues, respectively (Figure 5A, the second and third bars). In contrast, nc-circRNAs in both tissues decreased to 34.9%.

Beyond the circRNAs exclusively expressed in either tumor or tumor-adjacent tissues, many circRNAs that occurred in both tissues also exhibited differential expression. Indeed, 6 and 301 nc-circRNAs, as well as 11 and 1,210 c-circRNAs, of the total 23,028 shared circRNAs were statistically significantly up- and down- regulated (Supplemental Table S11), respectively, in tumors versus tumor-adjacent tissues (Figure 5C), consistent with early results from other circRNA pipelines^[66]^. Combined, these results indicated that nc- circRNAs, particularly the complete-nonboundary subtype, exhibit strong tissue-specificity.

To appreciate the potential involvement of circRNAs in NSCLC, we examined the functions enriched by the host genes of differentially expressed circRNAs. As only a few circRNAs were up-regulated in tumors, we selected tumor-specific circRNAs for analysis, defined as those present in at least 50% of tumor samples but absent in all tumor-adjacent tissues (Supplemental Table S11). Function enrichment analysis (GO) of the host genes of 1,283 tumor-specific circRNAs revealed significant associations with pathways related to cell migration, genomic stability/chromatin remodeling, transcriptional regulation, and cell signaling (Figure 5D; Supplemental Table S12), which are characteristic of cancers. Concordant pathway results were also observed on the host genes of tumor-adjacent specific circRNAs and tumor down-regulated circRNAs (Supplemental Figures S2B and S2C; Supplemental Table S12).

Although more circRNAs appeared in tumor-adjacent samples than in tumors (Figure 5B), a substantial number of circRNAs were exclusively expressed in all four tumor samples. The host genes of the top 110 tumor-exclusive circRNAs included PAK1, CDCA7, NUSAP1, BARD1, and CBLC, which have been previously implicated in tumor proliferation, invasion, and poor clinical outcomes in NSCLC. In particular, PAK1^[67–69]^, CDCA7^[70–72]^ and CBLC^[73–75]^ promote tumor growth, metastasis, and therapy resistance; NUSAP1^[76–78]^ and BARD1^[79, 80]^ are also known to be elevated in NSCLC and proposed as potential biomarkers or therapeutic targets. Similarly, tumor-upregulated circRNAs from the shared set were predominantly derived from genes with recognized roles in lung cancer, many of which have been reported to promote cancer development. These include three genes (USP4^[81–84]^, ASH1L^[85, 86]^, and COPG1^[87]^) that host c-circRNAs, and three (TOM1L1^[88–92]^, HOOK3^[93–95]^, and RAPGEF5^[96]^) that accommodate nc-circRNAs (Figure 5C). Importantly, circRNAs from ASH1L^[97]^, PRKAR1B^[98, 99]^, and RAPGEF5^[100]^ have been reported to be involved in cancer development, suggesting that they might also be involved in NSCLC.

In summary, all these results provided the first glimpse into the potential circRNA functions in NSCLC, suggesting their potential clinical relevance. The tumor-specificity of many circRNAs, particularly nc-circRNAs, underscores their potential roles in tumorigenesis and highlights the need for functional studies to elucidate their mechanisms of action in NSCLC.

## 3. DISCUSSION

This study significantly expanded our understanding of circular RNA biology and the clinical potential of circRNAs by systematically identifying and characterizing nc-circRNAs.

### 3.1. The CircRNA Landscape is More Complex and Diverse Than Anticipated

Our findings revealed, for the first time, that the circRNA landscape was far more complex and extensive than previously appreciated, with nc-circRNAs originating within exons, introns, and intergenic regions, constituting the majority (>75%) of circular transcripts in lung adenocarcinoma cells (Figure 1A) and NSCLC (Figure 5A). The higher average expression levels of nc-circRNAs compared to their canonical counterparts, combined with their evolutionary conservation across species, strongly argue against these molecules representing transcriptional noise or methodological artifacts. Our multi-assay experimental validation confirmed that nc-circRNAs were constitutive members of the circRNA family. Nevertheless, our sequence feature analysis revealed that nc-circRNAs, particularly the complete-nonboundary and intergenic subtypes were not directly associated with U2 or U12 signals. While this result indicates that nc-circRNAs may follow their own biogenesis or are products of some unexplored variants of splicing mechanisms, a topic for future research.

### 3.2. Noncanonical CircRNAs are Involved in Cell Cycle and Cancer-related Pathways in Lung Cancer

Our study provided the first experimental evidence that nc-circRNAs were functional molecules with potential clinical relevance in cancer. We investigated two prototypic nc-circRNAs: exonic nc-circDNAJBP1 and intergenic nc-circINT1. They play distinct roles in the proliferation and survival of lung adenocarcinoma cells.

Our investigation revealed that nc-circDNAJB6P1, which is primarily cytoplasmic (Figure 3D), is a potent suppressor of oncogenic phenotypes, particularly cell migration. This exonic circRNA originates from the pseudogene *DNAJB6P1*, a derivative of the *DNAJB6* gene, which is itself a known tumor and metastasis suppressor. Depleting nc-circDNAJB6P1 significantly enhances the migratory capacity of lung cancer cells (Figure 4E, 4F), providing a strong functional link between this pseudogene-derived circRNA and a critical hallmark of cancer progression, positioning it as a key regulator of metastasis. In addition, nc-circINT1, localized primarily to the nucleus (Figure 3D), influenced chromatin organization by modulating histone genes (HIST1H3B/G/H and H3F3A/C) (Figure 4L) and participated in DNA damage response via ATM/ATR regulation (Figure 4M). Its knockdown triggered an inflammatory response, characterized by elevated cytokine expression and activation of the JAK-STAT and MAPK pathways through IL-8 and LIF (Figure 4N). These findings positioned nc-circINT1 as a multifunctional regulator of chromatin dynamics, genome stability, and inflammatory signaling.

The effects of these nc-circRNAs highlighted their participation in complementary regulatory networks that fine-tune cellular homeostasis. Our findings expanded the functional repertoire of circRNAs, revealing their ability to influence critical cancer pathways through diverse mechanisms beyond microRNA sequestration. The involvement of nc-circINT1 in chromatin maintenance underscored previously unrecognized layers of circRNA-mediated gene regulation with implications for cancer biology and therapeutics. More broadly, our discovery that intergenic regions can produce functional circRNAs challenged conventional definitions of transcriptional units and expanded the functional landscape of the noncoding genome.

The PCR- and sequencing-based circRNA profiling in NSCLC revealed a substantial divergence in circRNA expression between tumors and tumor-adjacent tissues, in both abundance and subtype composition. Notably, nc-circRNAs, particularly those of the complete-nonboundary type, exhibited markedly higher tumor specificity than the canonical forms, suggesting their potential involvement in tumor-specific regulatory networks. Functional enrichment of differentially expressed circRNAs across tumors, tumor- adjacent tissues, and both samples revealed convergence on pathways related to epithelial-mesenchymal transition, chromatin dynamics, and tissue integrity, implicating circRNAs in both tumor progression and structural remodeling of lung tissues. The consistent detection of several tumor-specific circRNAs across all samples, with host genes known to drive oncogenesis, further supported their cancer relevance. These findings positioned nc-circRNAs as underappreciated gene regulators with functional and diagnostic potential in NSCLC, warranting future mechanistic studies to dissect their roles in disease initiation and progression.

### 3.3. Methodological Advance – The CAT Algorithm

Developing an annotation-independent method posed technical challenges due to the lack of exon-intron boundary information, thereby significantly expanding the pool of candidate BSJs. Alignment of a single sequencing read to multiple genomic loci, primarily due to low-complexity regions, pseudogenes, and paralogs, exacerbated the problem of determining the exact genomic origin of a circRNA.

Our novel CAT algorithm represented a significant methodological advance. It operates independently of genome annotation to detect both canonical and noncanonical circRNAs in an unbiased manner. Our extensive benchmark analysis showed that CAT performed comparably to leading methods for detecting canonical circRNAs (Figure 1D) and also effectively extended this capability to noncanonical classes. CAT’s performance is in part attributable to its stringent filtering criteria, which require multiple supporting RNA- seq reads covering regions adjacent to candidate BSJs.

Experimental validation confirmed CAT’s robust performance, achieving approximately 80% prediction accuracy for both canonical and noncanonical circRNAs. Notably, CAT identified 664 high-confidence canonical circRNAs that have been missed by all existing methods, overcoming inherent biases and advancing circRNA research.

### 3.4. Distinct Genomic Features Define Canonical and Noncanonical circRNAs

Our comprehensive analysis of circRNA origins reveals distinct genomic and sequence architectures that distinguish canonical and noncanonical circRNAs. Evaluation of splicing signals highlights a primary divergence between these two classes. Most (87%) canonical circRNAs are adjacent to major or minor splice sites (Supplemental Figures S1B), such splicing signals are almost completely absent in the intergenic and complete-nonboundary subtypes (Figure 1C). Furthermore, we identified a clear bias in the genomic origin of exonic nc-circRNAs. They are preferentially derived from genes with more compact architectures, characterized by fewer and shorter exons (Figures 2B, 2C, 2F).

Collectively, these findings establish a clear set of genomic and sequence hallmarks that differentiate circRNA subtypes: the divergent reliance on splicing signals and the distinct genomic architectures of their host genes. These features not only underscore the unanticipated diversity of RNA circularization mechanisms but also provide a critical foundation for future mechanistic investigations into their biogenesis. This new finding has immediate practical implications for improving the accuracy of circRNA prediction algorithms and for the functional interpretation of these molecules in health and disease.

In summary, our study not only advances the methodological frontier of circRNA identification but also deepens our understanding of the diversity and complexity of circRNA genomic origins and cellular functions, opening new avenues for research into their functional and regulatory roles and clinical significance.

## 4. MATERIALS and METHODS

### 4.1. Cell-based Experiments

#### 4.1.1. Sample collection and RNA sequencing

Cell lines NCI-H23, A549, and HSAEC were cultured in RPMI 1640 Medium (Gibco), Ham’s F-12K (Kaighn’s) Medium (Gibco), and Bronchial Epithelial Cell Growth Medium (ATCC), respectively, supplemented with 10% Fetal Bovine Serum (FBS) (Gibco) and 1% penicillin/streptomycin (P/S) (Gibco) in a humidified 5% CO_2_ incubator at 37°C. Total RNA was extracted using Trizol (ThermoFisher) according to the manufacturer’s instructions. RNA integrity and concentration were analyzed using a uDrop Duo Plate in a Multiskan SkyHigh Microplate Reader (ThermoFisher). Reverse transcription was performed with SuperScript IV First-Strand Synthesis System (ThermoFisher).

For RNA-seq library preparation, the NEBNext Ultra II DNA Library Prep Kit for Illumina (New England Biolabs) was used. The library was sequenced on a NovoSeq 6000 instrument using a Mid Output Kit v2.5 (150 cycles) (Illumina), generating approximately 50 million paired-end 150-nucleotide reads per library.

#### 4.1.2. Validation

***PCR and qPCR*** were performed using divergent primers designed to span across the BSJs of the circRNAs to be analyzed; correspondingly, convergent primers were also included for linear RNAs as negative controls (Figure 3A, top panel). PCR was performed using complementary DNA (cDNA) and genomic DNA (gDNA) templates to verify the presence of circRNAs derived specifically from RNA, rather than from genomic DNA or contamination.

***Sanger sequencing***, conducted by Sangon Biotech, involved inserting a T vector into purified PCR products. The full lengths of the circRNA candidates were determined, and the junction sites were confirmed using the constructed primers (WZ Biosciences Lnc) listed in Supplemental Table S3.

***Amplicon sequencing*** was performed by Novogene using an amplicon pool containing 2 μL of PCR products from the circRNA candidates to be analyzed (see Table S3 for the PCR primers). The PCR products were cleaned using Amicon Ultra Centrifugal Filter, 10kDa MWCO columns (MIllipore). The concentration was assessed using the uDrop Duo Plate in a Multiskan Sky High Microplate Reader (ThermoFisher). For details on the analysis of amplicon sequencing results, see the section in the Supplemental Methods: Amplicon Sequencing Analysis.

***RNase R treatment*** followed by PCR/qPCR. Following total RNA extraction, 234 μg of total RNA was incubated for 10 minutes at 37°C with 2 U/μg RNase R (Beyotime Biotechnology) added to the treated group to enrich circRNAs and degrade linear RNAs. CircRNA expression abundances were analyzed by comparing circRNA samples to an equivalent amount of total RNA without RNase R treatment, using quantitative PCR with SYBR Green qPCR Master Mix (QIAGEN).

#### 4.1.3. Cell fraction assay

The cytoplasmic and nuclear fractions were separated and isolated from cells using a PARIS kit (Life Technologies) according to the manufacturer’s instructions. Briefly, chondrocytes were lysed with a cell fractionation buffer and centrifuged. The supernatant was then transferred to new RNase-free EP tubes, and the remaining lysate was washed with cell fractionation buffer and centrifuged. The nuclei were lysed with a cell disruption buffer. The resulting lysate and supernatant were mixed with a 2×lysis binding solution, and an equal volume of ethanol was added through a filter cartridge. The sample was subsequently washed with a series of wash solutions. Finally, RNAs from the cytoplasm and nuclei were eluted using the elution solution. Cytoplasmic and nuclear RNAs were reverse-transcribed into cDNAs and detected and analyzed by qPCR.

#### 4.1.4. Function analysis

##### Cell proliferation assay

Cell viability was assessed using the Cell Counting Kit-8 (CCK-8) assay. The NCI-H23 and A549 cells were seeded at a density of 5 × 10^4^ cells per well in a 24-well plate and incubated at 37°C. After overnight culturing, the cells were transfected with siRNA and incubated for 24 h. Then, 100 μL CCK-8 reagent (MCE) was added to the culture medium and incubated at 37°C for 2 h. The absorbance was measured at 450 nm using a microplate reader (Thermo Fisher Scientific).

##### Cell apoptosis

Cell apoptosis was measured by flow cytometry. The H23 cell was seeded at a density of 4 × 10^5^ cells per well in a 6-well plate and incubated at 37°C. After overnight culturing, the cells were transfected with siRNA and incubated for 24 h. After cell digestion, cells were collected, washed with PBS (Gibco) and filtered through a 40-mm cell strainer (Thermo Fisher Scientific). The cells were stained with PI/Hoechst 33342 (Solarbio). After incubation for 30 min at 4 ℃ with no light, cell apoptosis was detected using flow cytometry (BD FACSymphony A3 Cell Analyzer).

##### Wound Healing Assay

Cell migration was assessed using a wound-healing assay. A549 cells were seeded at 2.5 × 10⁵ cells per well in 6-well plates and grown for 24 h to form a confluent monolayer. Following transfection with siRNAs and a further 24 h incubation, a linear scratch wound was generated in the center of each monolayer using a sterile 1000 µL pipette tip. Cellular debris was removed by washing with PBS, and the wells were replenished with fresh complete medium containing 10% FBS. Images of the wound were captured at 0, 24, and 48 h using an inverted microscope. The rate of wound closure was quantified by measuring the change in wound width over time.

##### Transwell migration assay

Transwell migration assays (Corning, NY, USA) were performed to further quantify cell migratory capacity. A549 cells were prepared as described above (seeded, grown for 24 h, and transfected for 24 h). Post-transfection, the cells were harvested and resuspended at 5 × 10⁵ cells/mL in serum-free medium. A 150 µL aliquot of the cell suspension was then added to the upper chamber of a Transwell insert. The lower chamber was filled with 700 µL of medium containing 30% FBS, which served as a chemoattractant. After 24 h of incubation at 37 °C, non-migrated cells remaining on the upper surface of the membrane were carefully removed with a cotton swab. Cells that had migrated to the lower surface were fixed in 4% paraformaldehyde and stained with crystal violet. The number of migrated cells was determined by counting five randomly selected fields per insert under a microscope.

##### NanoString

Cultured H23 after RNAi was followed by RNA extraction using TRIzol (ThermoFisher) according to the manufacturer’s protocol. The RNA concentration was determined by using a microplate reader. Then, multiplex gene expression analysis was performed using the NanoString nCounter® Technology with the Mouse Neuroinflammation Panel and nCounter® SPRINT™ Profiler according to the manufacturer’s protocol (NanoString Technologies, USA)^[101]^. The data were analyzed using nSolver 4.0 software. Our analysis included background subtraction using Negative Controls, standard normalization using Positive Control Normalization, and CodeSet Content Normalization.

##### Human participants

Patients with non-small cell lung cancer (NSCLC) were recruited from the affiliated hospital of Xiamen University School of Medicine. All four participants provided written informed consent after receiving detailed study descriptions. The study was approved by the hospital’s Ethics Committee (Institutional Review Board).

### 4.2. Computational Analyses

Some details of the CAT method and data analysis were documented in the Supplemental Methods file.

#### 4.2.1. The CAT pipeline for identifying Circular RNA of All Types

CAT employed a systematic approach, relying solely on RNA-seq data without genome annotation, to comprehensively identify all circRNAs present in the genome as captured by the given RNA-seq data. It utilized strand-specific mapping to detect circRNAs originating from both strands of the genome. The modular design of the CAT pipeline accommodates data from both single-end and paired-end sequencing formats. Additionally, by integrating UMI (unique molecular identifier) and barcode information, CAT can identify and profile circRNAs at single-cell resolution using single-cell RNA-seq data.

The CAT pipeline first mapped RNA-seq reads to the reference genome using STAR^[102]^ (Figure 1B, Step 1), retaining unmapped reads for further analysis for finding candidate BSJs. To detect BSJs, CAT extracted short sequences, called anchors, from both ends of each unmapped read and aligned them to the reference genome (Figure 1B, Step 2). The initial length of the anchors was set to 1/5 of the read length, up to 40 bp, to optimize efficiency. Successfully mapped anchor pairs underwent an extension process in which the anchors were progressively extended toward each other along the read sequence while maintaining alignment with their respective genomic locations. This extension process continued until either a BSJ was detected or the number of mismatches exceeded a length-dependent threshold of l = ⌊L/100⌋ for reads with L nucleotides.

To reduce the likelihood of false-positive BSJs caused by mismatches and multi-locus mapping during read alignment to the genome, a filter based on linear supporting reads was introduced (Figure 1B, Step 3). A BSJ was considered valid only if there were reads that were mapped to both the upstream and downstream 5 bp long genomic regions adjacent to the BSJ, with read coverages of the adjacent regions obtained from the mapping result of step 1. Finally, the number of reads aligned to a BSJ was used as a proxy for the expression of the corresponding circRNAs at the BSJs. To accurately calculate this electronic expression, all unmapped reads from Step 1 were realigned to this junction, and we reported only circRNAs with at least 5 supporting reads covering the junction (Figure 1B, Step 3).

A narrative comparison of the CAT method with the existing methods for finding canonical circRNAs (listed in Supplemental Table S1) is warranted. These methods are thoroughly compared and analyzed in the latest comparative study.^[19]^ In brief, these methods consist of two primary steps: detecting candidate BSJs and subsequently filtering out potential spurious junctions. Depending on the specific strategies adopted, they can be grouped into three categories: pseudo-reference-based, segmented-read-based, and integrative approaches.^[103]^ Leveraging the back-splicing biogenesis model, the pseudo-reference-based approach uses genome annotation to construct possible BSJ sequences from all combinations of annotated exons across all genes. Junctions constructed from RNA-seq reads that align with RNA-seq reads are taken as candidate BSJs. The segmented-read-based approach relies on splitting reads that fail to map to the genome in the first place, allowing their split segments to be aligned to the genome separately. This enables the splitting points to help locate BSJs (Figure 1B). Although methods following this strategy, in principle, can detect candidate BSJs across the genome, they subsequently filter out many of them based on the genome annotation or splicing signals. In other words, these methods use genome annotation to determine whether candidate BSJs are genuine. Lastly, the integrative approach combines several existing methods, thereby inherently magnifying their drawbacks.

The CAT method, in principle, falls into the category of the segmented-read-based approach. It deviates from the existing method in the parameters used, such as the number of mismatches allowed and how to handle multiple mapping results, and more critically, in the filtering step. CAT relies completely on sequencing reads rather than genome annotations to filter out spurious back-splicing junctions. Thus, it can identify all types of circRNA with diverse genomic origins. For details on the implementation and parameter settings of CAT, see the section in the Supplemental Methods: Circular RNA Detection Using the CAT Pipeline.

#### 4.2.2. Classification and annotation of circular RNA of all types

Although CAT did not rely on genome annotation to detect circRNAs, it utilized the human genome annotation (version GRCh38) in a strictly strand-specific manner to stratify them into different classes, thereby helping to appreciate their diverse genomic origins (Figure 1A) and potential functions. The first class consists of c-circRNAs with both BSJs aligning with exon-intron boundaries of protein-coding genes or long noncoding RNAs (lncRNAs) with exon-intron structures. The second class is complete noncanonical circRNAs with none of their BSJs matching any annotated exon-intron boundaries. The third class is half boundary circRNAs with one, but not both, of their BSJs mapping to exon-intron boundaries. Note that lariat circRNAs have similar structures to half-boudary circRNAs, so they were classified in a separate category. CircRNAs in these four classes, except for some from lncRNAs, are genic. The fifth class is intergenic, consisting of circRNAs with at least one BSJ in regions devoid of gene annotation on the transcribed strand. Importantly, owing to this strict strand-specific criterion, antisense circRNAs—which lack exon-intron annotations on their respective strand despite overlapping with annotated genes on the opposite strand—are categorized into this intergenic class.

#### 4.2.3. Performance assessment

CAT’s performance was evaluated on the benchmarking dataset from NCI-H23 lung cancer cells, which produced 309,485,791 strand-specific paired-end reads (151 bp) and 234 c-circRNAs validated by three orthogonal methods: qPCR, RNase R treatment, and amplicon sequencing as done in the benchmarking study.^[19]^ The sensitivity for detecting circRNAs was calculated as the proportion of these validated circRNAs that were successfully detected. The precision compared the identified circRNAs by the method against those in the 13 public circRNA databases, including Circ2Disease, circad, CircAtlas, circbank, circBase, CIRCpediav2, CircR2disease, CircRiC, circRNADb, CSCD, exoRBase, MiOncoCirc and TSCD^[41–52, 104]^. Notably, CAT’s precision assessment excluded noncanonical circRNAs, as these are largely absent from the existing databases.

#### 4.2.4. Statistical analysis

All statistical analyses were performed using GraphPad Prism software. Statistical significance between groups was determined using an unpaired two-tailed Student’s t-test, with p < 0.05 as the significance threshold.

#### 4.2.5. Classification of U2- and U12-type spliceosome signals

To systematically investigate the potential involvement of the minor spliceosome in circRNA biogenesis, we analyzed the splice signals at the BSJs using reference data from the Intron Annotation and Orthology Database (IAOD).^[55]^ Reference genomic coordinates for U2-type (major) and U12-type (minor) intron boundaries were obtained from the IAOD. After identifying BSJs, the algorithm extracted the exact genomic coordinates of the junction sites from both ends. To account for potential positional ambiguity at the junction sites, a strict ±1 bp tolerance window was applied to expand these junction regions. The algorithm then intersected these expanded regions with the annotated coordinates of U2 and U12 intron boundaries. Based on whether any reference intron boundary position fell within the expanded regions, each circRNA was classified into one of four categories: containing U2 signals, containing U12 signals, containing both, or neither. This precise region-based matching enabled the quantification of canonical and noncanonical spliceosome contributions across all circRNA subtypes.

#### 4.2.6. Function enrichment analyses

Functional characterization was performed by integrating KEGG and GO analyses using the GSEApy Python module with Enrichr API (http://amp.pharm.mssm.edu/Enrichr). Enrichment analysis employed KEGG_2021_Human and GO_Biological_Process_2023 databases, filtered by adjusted *p*-value threshold of 0.05.

#### 4.2.7. Coding potential analysis

The assessment of circRNAs’ coding capacity used ribosome profiling data from A549 cells^[61]^ and internal Ribosome entry site (IRES) prediction. The capture of BSJ sequences by the Ribosome indicates that the corresponding circRNAs can be loaded into the Ribosome complex and may be subsequently translated into proteins. A549 ribo-seq data were aligned to the BSJ sequences from A549 cells using bowtie2, and we matched BSJ sequences from A549 cells with IRES references from the IRESbase database^[63]^ using BLASTN (≥80% identity, ≥30 nucleotides). Furthermore, open reading frame (ORF) analysis was applied to sequences between BSJs to assess their protein-coding potential, accounting for rolling-circle translation via fourfold sequence multiplication.^[62]^ All ORFs beginning with an AUG initiation codon were identified separately for each circRNA and further filtered based on the requirements of a minimum length of 20 amino acids and spanning the corresponding BSJs.^[64]^ The subsections in the Supplemental Methods, specifically “Ribosome Profiling Analysis”and “Internal Ribosome Entry Site Analysis,”provide detailed descriptions of the analysis process and specific parameters related to coding potential.

#### 4.2.8. Single-cell RNA-seq data analysis

We analyzed a random primer library dataset from mixed cancer cell lines (A549, HCC827, and K562). Raw sequencing data were processed using SeekSoulTools with GRCh38 as the reference. Quality control measures included filtering based on gene count, UMI count, mitochondrial gene percentage, and doublet removal using Scrublet to obtain high-quality single cells. The gene expression matrix was analyzed using the Scanpy pipeline for normalization, dimensionality reduction, and Leiden clustering. Uniform Manifold Approximation and Projection (UMAP) was employed for visualization. Cell types were annotated based on established marker genes for both the cell line mixture (see the section in Supplemental Methods: Single- cell processing for detailed parameters and marker genes).

## DATA AVAILABILITY

The CAT pipeline and figure-drawing scripts are available on GitHub at https://github.com/GenomicMedicine/CAT.

The circRNAs listed in databases (Circ2Disease, circad, CircAtlas, circbank, circBase, CIRCpediav2, CircR2disease, CircRiC, circRNADb, CSCD, exoRBase, MiOncoCirc and TSCD) can be accessed from a recent comparative study^[19]^.

Public NCI-H23 cell line RNA datasets used in the study: SRX13414573 (untreated NCI-H23), SRX13414576 (RNase R-treated NCI-H23). Public A549 Ribo-seq: GSE133111. Public single-cell mixed cancer cell lines can be accessed at http://seekgene-public.oss-cn-beijing.aliyuncs.com/software/data/demodata/cellline.tar.gz. Public RNA-seq datasets of paired tumor and tumor-adjacent tissues are available in the NCBI BioProject database under accession number PRJNA863919.

The in-house RNA-seq data from NCI-H23, A549, and HSAEC cells generated in this study: The raw sequence data have been deposited in the Genome Sequence Archive^[105]^ of the National Genomics Data Center,^[106]^ China National Center for Bioinformation/Beijing Institute of Genomics, Chinese Academy of Sciences, accessible at https://ngdc.cncb.ac.cn/gsa under access number GSA:HRA010971.

## SUPPLEMENTARY DATA

Supplemental Methods; Supplemental Tables 1-11; Supplemental Figures 1-2.

## DECLARATION OF GENERATIVE AI AND AI-ASSISTED TECHNOLOGIES IN THE WRITING PROCESS

During manuscript preparation, the authors used Claude or Perplexity to refine the language in some of their original writing and subsequently reviewed and edited the content as needed. The authors take full responsibility for the content of the work.

## AUTHOR CONTRIBUTIONS

WZ and SQ conceived, designed, and supervised the research. WZ, KL and WW designed the CAT method, and KL and WW implemented the CAT pipeline. SQ and WZ designed molecular experiments, and AEN, YZ, JD, HQ and JZ prepared the RNA-seq libraries and performed the validation and functional experiments. KM analyzed the single-cell data. PW, DL, F-M K, and WCC provided reagents. SQ, LK and WZ analyzed the data and results. WZ, SQ and LK wrote and revised the manuscript.

## ACKNOWLEDGMENTS

The work was supported in part by funding from the Hong Kong RGC theme-based Strategic Target Grant Scheme (STG STG1/M-501/23-N), NSFC/RGC Collaborative Research Scheme (CRS_HKBU 2021/22), the Hong Kong RGC Collaborative Research Fund (CRF C5005-23WF), the Hong Kong Global STEM Professor Scheme, and the Hong Kong Jockey Club Charities Trust.

---

**Figure S1.**
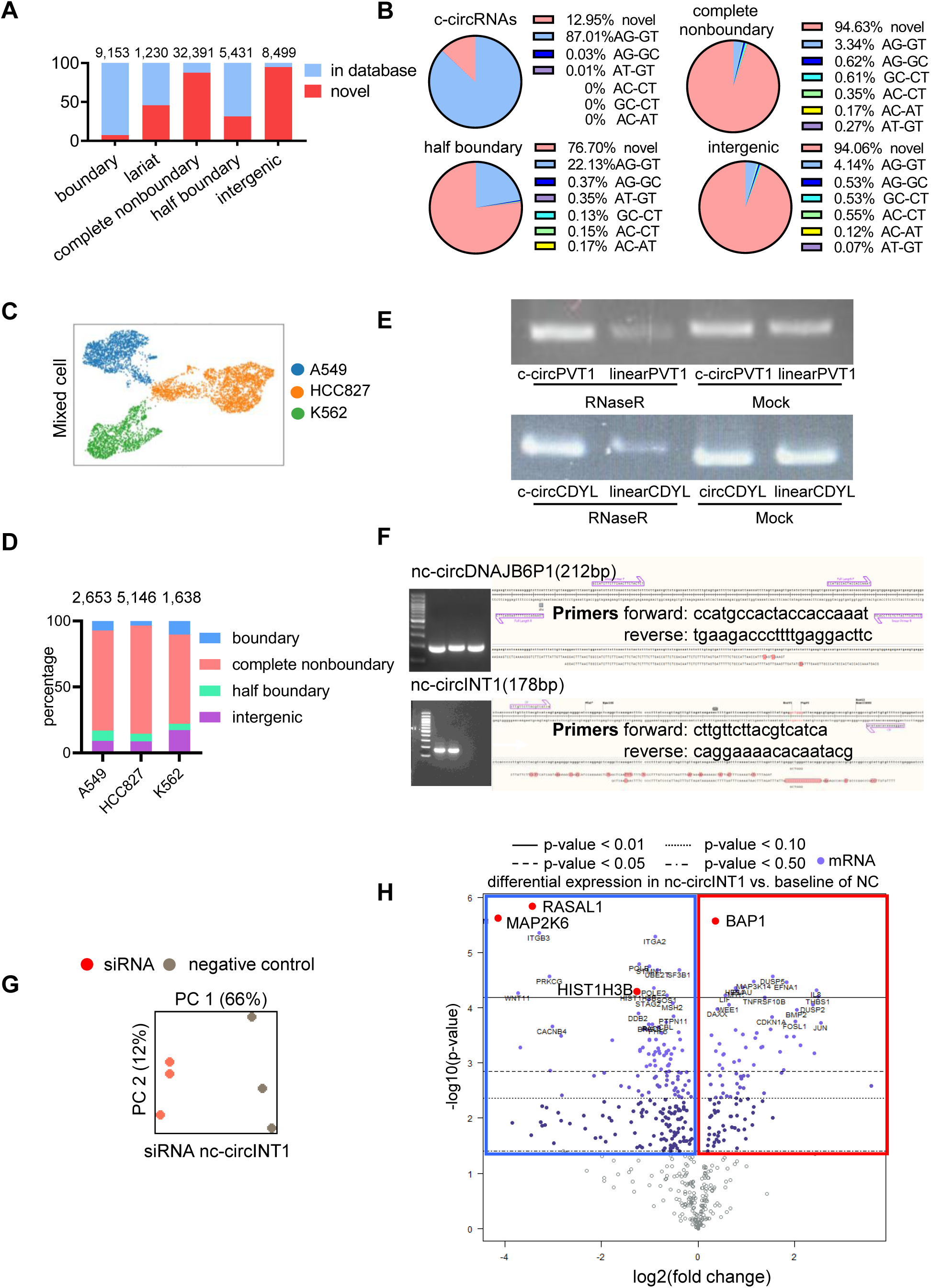
Characterization of circRNA function and sample information. (A) CircRNA landscape of NCI-H23 cells. The stacked bar chart illustrates the proportions of novel and known circRNAs for each category. (B) The types and distributions of known and unknown splice signals at BSJs of different subtypes of circRNAs. (C) Application of CAT to single-cell RNA-seq data to identify circRNAs in single-cell resolution. UMAP visualization of cell clustering based on marker gene expression in mixed cancer cell lines. (D) Distributions of different circRNA types in single cells. Shown are the relative abundances of different circRNA subtypes in cell lines. (E) The agarose gel electrophoresis results of the PCR products of two canonical circRNAs. (F) Sanger sequencing results of nc-circDNAJB6P1 and nc-circINT1. (G) Principal Component Analysis (PCA) of gene expression profiles following nc-circINT1 siRNA-mediated knockdown. PC1 and PC2 values shown in parentheses are the percentages of variance explained. (H) Volcano plots showing differential gene expression after nc-circINT1 knockdown. The x-axis is in log2(fold change) and the y-axis in −log10(*p*-value). Red dots correspond to those genes with important functions and significantly differential expression. Blue and red quadrants indicate down- and up-regulated genes, respectively.

**Figure S2.**
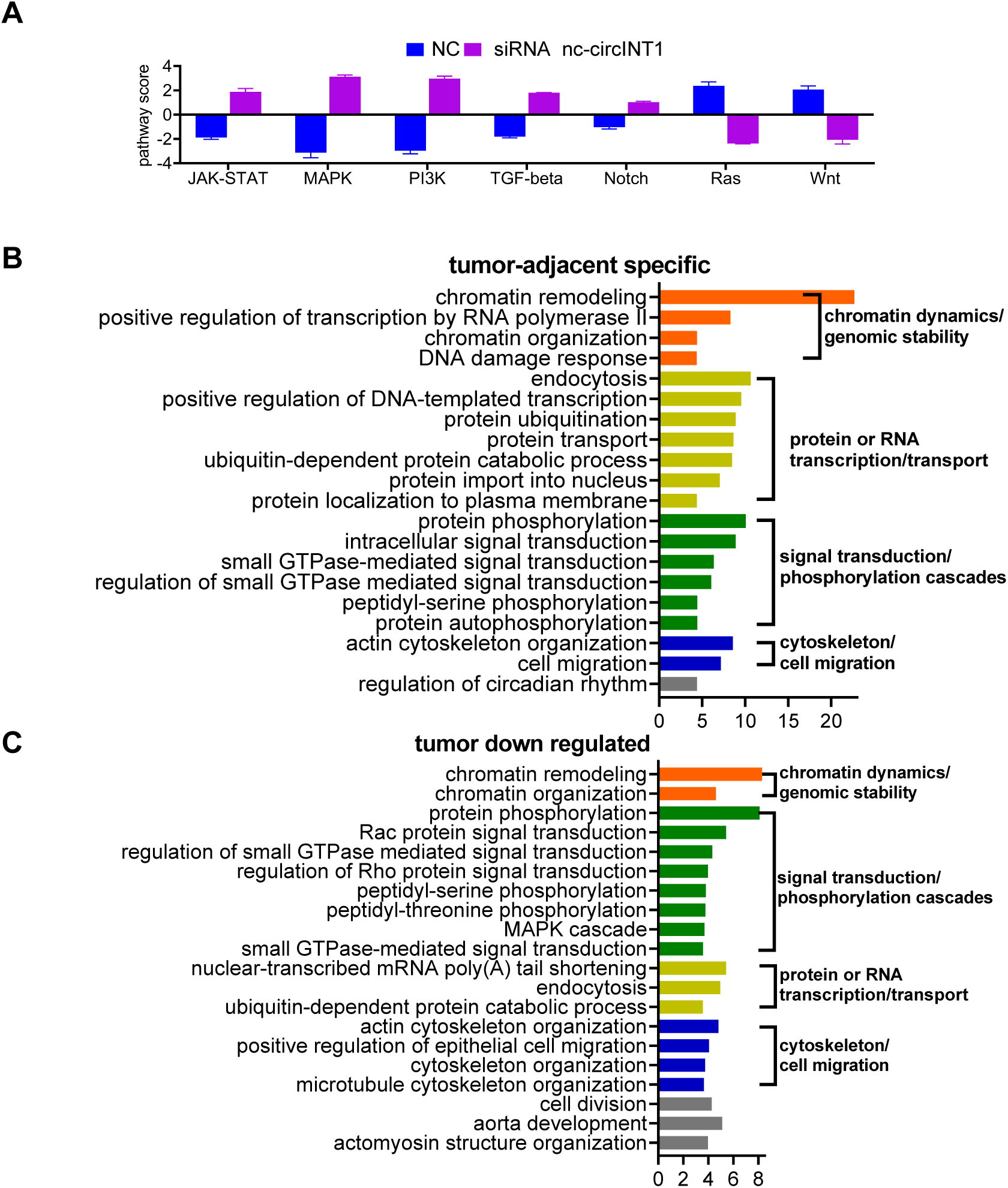
Signaling alterations following siRNA-mediated knockdown in tumor cells and enrichment analysis of tumor-specific circRNAs. (A) Pathway activity analysis reveals changes in major signaling pathways after knockdown of nc-circINT1. Bars represent pathway scores relative to negative control (NC). Positive and negative values indicate pathway activation and suppression, respectively. (B) GO enrichment analysis of host genes from tumor-adjacent specific circRNAs. (C) GO enrichment analysis of host genes from tumor down-regulated circRNAs.

## REFERENCES

1. Memczak, S., Circular RNAs are a large class of animal RNAs with regulatory potency. Nature, 2013. 495.

2. Hansen, T.B., et al., Natural RNA circles function as efficient microRNA sponges. Nature, 2013. 495(7441): p. 384–8.

3. Liu, C.X. and L.L. Chen, Circular RNAs: Characterization, cellular roles, and applications. Cell, 2022. 185(12): p. 2016–2034.

4. Hsu, M.-T. and M. Coca-Prados, Electron microscopic evidence for the circular form of RNA in the cytoplasm of eukaryotic cells. Nature, 1979. 280(5720): p. 339–340.

5. Salzman, J., et al., Circular RNAs Are the Predominant Transcript Isoform from Hundreds of Human Genes in Diverse Cell Types. Plos One, 2012. 7(2).

6. Li, Z., et al., Exon-intron circular RNAs regulate transcription in the nucleus. Nat Struct Mol Biol, 2015. 22(3): p. 256–64.

7. Ashwal-Fluss, R., et al., circRNA biogenesis competes with pre-mRNA splicing. Molecular cell, 2014. 56(1): p. 55–66.

8. Huang, D., et al., Tumour circular RNAs elicit anti-tumour immunity by encoding cryptic peptides. Nature, 2024. 625(7995): p. 593–602.

9. Yu, X., et al., Circular RNAs: New players involved in the regulation of cognition and cognitive diseases. Front Neurosci, 2023. 17: p. 1097878.

10. Joaquim, V.H.A., et al., Circular RNAs as a Diagnostic and Therapeutic Target in Cardiovascular Diseases. Int J Mol Sci, 2023. 24(3).

11. Vo, J.N., et al., The Landscape of Circular RNA in Cancer. Cell, 2019. 176(4): p. 869–881 e13.

12. Conn, V.M., A.M. Chinnaiyan, and S.J. Conn, Circular RNA in cancer. Nature Reviews Cancer, 2024. 24(9): p. 597–613.

13. Ghafouri-Fard, S., et al., A concise review on the role of CircPVT1 in tumorigenesis, drug sensitivity, and cancer prognosis. Frontiers in Oncology, 2021. 11: p. 762960.

14. Zheng, F. and R. Xu, CircPVT1 contributes to chemotherapy resistance of lung adenocarcinoma through miR-145-5p/ABCC1 axis. Biomedicine & Pharmacotherapy, 2020. 124: p. 109828.

15. Chen, I., C.Y. Chen, and T.J. Chuang, Biogenesis, identification, and function of exonic circular RNAs. Wiley Interdiscip Rev RNA, 2015. 6(5): p. 563–79.

16. Zhong, Y., et al., Systematic identification and characterization of exon-intron circRNAs. Genome Res, 2024. 34(3): p. 376–393.

17. Chen, L.L., The biogenesis and emerging roles of circular RNAs. Nat Rev Mol Cell Biol, 2016. 17(4): p. 205–11.

18. Li, X., et al., The mechanism and detection of alternative splicing events in circular RNAs. PeerJ, 2020. 8: p. e10032.

19. Vromman, M., et al., Large-scale benchmarking of circRNA detection tools reveals large differences in sensitivity but not in precision. Nat Methods, 2023. 20(8): p. 1159–1169.

20. Liu, X., et al., Interior circular RNA. RNA Biol, 2020. 17(1): p. 87–97.

21. Guarnerio, J., et al., Oncogenic Role of Fusion-circRNAs Derived from Cancer-Associated Chromosomal Translocations. Cell, 2016. 165(2): p. 289–302.

22. Zhang, Y., et al., Circular intronic long noncoding RNAs. Mol Cell, 2013. 51(6): p. 792–806.

23. Robic, A. and C. Kuhn, Beyond Back Splicing, a Still Poorly Explored World: Non-Canonical Circular RNAs. Genes (Basel), 2020. 11(9).

24. Ye, C.Y., et al., Widespread noncoding circular RNAs in plants. New Phytologist, 2015. 208(1): p. 88–95.

25. Gao, Y., J.F. Wang, and F.Q. Zhao, CIRI: an efficient and unbiased algorithm for de novo circular RNA identification. Genome Biology, 2015. 16.

26. Liu, X., et al., Canonical and Interior Circular RNAs Function as Competing Endogenous RNAs in Psoriatic Skin. Int J Mol Sci, 2021. 22(10).

27. Gao, Y., J. Zhang, and F. Zhao, Circular RNA identification based on multiple seed matching. Brief Bioinform, 2018. 19(5): p. 803–810.

28. Gaffo, E., et al., Sensitive, reliable and robust circRNA detection from RNA-seq with CirComPara2. Brief Bioinform, 2022. 23(1).

29. Chen, C.Y. and T.J. Chuang, NCLcomparator: systematically post-screening non-co-linear transcripts (circular, trans-spliced, or fusion RNAs) identified from various detectors. BMC Bioinformatics, 2019. 20(1): p. 3.

30. Chuang, T.J., NCLscan: accurate identification of non-co-linear transcripts (fusion, trans-splicing and circular RNA) with a good balance between sensitivity and precision. Nucleic Acids Res., 2016. 44.

31. Feng, J., Genome-wide identification of cancer-specific alternative splicing in circRNA. Mol. Cancer, 2019. 18.

32. Hoffmann, S., A multi-split mapping algorithm for circular RNA, splicing, trans-splicing and fusion detection. Genome Biol., 2014. 15.

33. Izuogu, O.G., Analysis of human ES cell differentiation establishes that the dominant isoforms of the lncRNAs RMST and FIRRE are circular. BMC Genomics, 2018. 19.

34. Jakobi, T., A. Uvarovskii, and C. Dieterich, Circtools: a one-stop software solution for circular RNA research. Bioinformatics, 2019. 35.

35. Li, M., Quantifying circular RNA expression from RNA-seq data using model-based framework. Bioinformatics, 2017. 33.

36. Ma, X.K., CIRCexplorer3: a CLEAR pipeline for direct comparison of circular and linear RNA expression. Genomics Proteomics Bioinformatics, 2019. 17.

37. Szabo, L., Statistically based splicing detection reveals neural enrichment and tissue-specific induction of circular RNA during human fetal development. Genome Biol., 2015. 16.

38. Westholm, J.O., Genome-wide analysis of Drosophila circular RNAs reveals their structural and sequence properties and age-dependent neural accumulation. Cell Rep., 2014. 9.

39. Ye, C.Y., Full-length sequence assembly reveals circular RNAs with diverse non-GT/AG splicing signals in rice. RNA Biol., 2017. 14.

40. Zhang, J., et al., Accurate quantification of circular RNAs identifies extensive circular isoform switching events. Nat. Commun., 2020. 11.

41. Chen, X.P., et al., circRNADb: A comprehensive database for human circular RNAs with protein-coding annotations. Scientific Reports, 2016. 6.

42. Dong, R., et al., CIRCpedia v2: An Updated Database for Comprehensive Circular RNA Annotation and Expression Comparison. Genomics Proteomics & Bioinformatics, 2018. 16(4): p. 226–233.

43. Fan, C.Y., et al., CircR2Disease: a manually curated database for experimentally supported circular RNAs associated with various diseases. Database-the Journal of Biological Databases and Curation, 2018.

44. Glazar, P., P. Papavasileiou, and N. Rajewsky, circBase: a database for circular RNAs. Rna, 2014. 20(11): p. 1666–1670.

45. Ji, P.F., et al., Expanded Expression Landscape and Prioritization of Circular RNAs in Mammals. Cell Reports, 2019. 26(12): p. 3444-+.

46. Li, S.L., et al., exoRBase: a database of circRNA, lncRNA and mRNA in human blood exosomes. Nucleic Acids Research, 2018. 46(D1): p. D106–D112.

47. Liu, M., et al., Circbank: a comprehensive database for circRNA with standard nomenclature. Rna Biology, 2019. 16(7): p. 899–905.

48. Rophina, M., et al., Circad: a comprehensive manually curated resource of circular RNA associated with diseases. Database, 2020. 2020: p. baaa019.

49. Ruan, H., et al., Comprehensive characterization of circular RNAs in ∼ 1000 human cancer cell lines. Genome Medicine, 2019. 11(1).

50. Xia, S.Y., et al., CSCD: a database for cancer-specific circular RNAs. Nucleic Acids Research, 2018. 46(D1): p. D925–D929.

51. Xia, S.Y., et al., Comprehensive characterization of tissue-specific circular RNAs in the human and mouse genomes. Briefings in Bioinformatics, 2017. 18(6): p. 984–992.

52. Yao, D.X., et al., Circ2Disease: a manually curated database of experimentally validated circRNAs in human disease. Scientific Reports, 2018. 8.

53. Wu, W., F. Zhao, and J. Zhang, circAtlas 3.0: a gateway to 3 million curated vertebrate circular RNAs based on a standardized nomenclature scheme. Nucleic Acids Research, 2024. 52(D1): p. D52–D60.

54. Chen, L.-L., et al., A guide to naming eukaryotic circular RNAs. Nature Cell Biology, 2023. 25(1): p. 1–5.

55. Moyer, D.C., et al., Comprehensive database and evolutionary dynamics of U12-type introns. Nucleic Acids Research, 2020. 48(13): p. 7066–7078.

56. Meyerhans, A., J.-P. Vartanian, and S. Wain-Hobson, DNA recombination during PCR. Nucleic acids research, 1990. 18(7): p. 1687–1691.

57. Chakravarti, D. and P.C. Mailander, Formation of template-switching artifacts by linear amplification. Journal of biomolecular techniques: JBT, 2008. 19(3): p. 184.

58. Li, J., et al., CircRM: profiling circular RNA modifications from nanopore direct RNA sequencing. Briefings in Bioinformatics, 2026. 27(1): p. bbaf726.

59. Wen, S.-y., J. Qadir, and B.B. Yang, Circular RNA translation: novel protein isoforms and clinical significance. Trends in molecular medicine, 2022. 28(5): p. 405–420.

60. Wu, P., et al., Emerging role of tumor-related functional peptides encoded by lncRNA and circRNA. Molecular cancer, 2020. 19: p. 1–14.

61. Campos, R.K., et al., Ribosomal stalk proteins RPLP1 and RPLP2 promote biogenesis of flaviviral and cellular multi-pass transmembrane proteins. Nucleic acids research, 2020. 48(17): p. 9872–9885.

62. Pamudurti, N.R., et al., Translation of circRNAs. Molecular cell, 2017. 66(1): p. 9–21. e7.

63. Zhao, J., et al., IRESbase: a comprehensive database of experimentally validated internal ribosome entry sites. Genomics, Proteomics and Bioinformatics, 2020. 18(2): p. 129–139.

64. Li, H., et al., riboCIRC: a comprehensive database of translatable circRNAs. Genome Biol, 2021. 22(1): p. 79.

65. Wu, L., et al., The E2F1–3 transcription factors are essential for cellular proliferation. Nature, 2001. 414(6862): p. 457–462.

66. Wang, Z.-H., et al., Circular RNA circFBXO7 attenuates non-small cell lung cancer tumorigenesis by sponging miR-296-3p to facilitate KLF15-mediated transcriptional activation of CDKN1A. Translational Oncology, 2023. 30: p. 101635.

67. Kim, E., et al., PAK1 tyrosine phosphorylation is required to induce epithelial-mesenchymal transition and radioresistance in lung cancer cells. Cancer Res, 2014. 74(19): p. 5520–31.

68. Chen, M.J., et al., PAK1 confers chemoresistance and poor outcome in non-small cell lung cancer via beta-catenin-mediated stemness. Sci Rep, 2016. 6: p. 34933.

69. Bu, H., et al., Therapeutic potential of IBP as an autophagy inducer for treating lung cancer via blocking PAK1/Akt/mTOR signaling. Mol Ther Oncolytics, 2021. 20: p. 82–93.

70. Li, S., et al., High expression of CDCA7 predicts tumor progression and poor prognosis in human colorectal cancer. Mol Med Rep, 2020. 22(1): p. 57–66.

71. Zhao, G., et al., Rare coexistence of three novel CDCA7-ALK, FSIP2-ALK, ALK-ERLEC1 fusions in a lung adenocarcinoma patient who responded to Crizotinib. Lung Cancer, 2021. 152: p. 189–192.

72. Zheng, D., et al., CDCA7 enhances STAT3 transcriptional activity to regulate aerobic glycolysis and promote pancreatic cancer progression and gemcitabine resistance. Cell Death Dis, 2025. 16(1): p. 68.

73. Li, Z., et al., CBLC promotes the development of colorectal cancer by promoting ABI1 degradation to activate the ERK signaling pathway. Transl Oncol, 2024. 45: p. 101992.

74. Hong, S.Y., et al., Stabilization of AURKA by the E3 ubiquitin ligase CBLC in lung adenocarcinoma. Oncogene, 2022. 41(13): p. 1907–1917.

75. Hong, S.Y., et al., Upregulation of E3 Ubiquitin Ligase CBLC Enhances EGFR Dysregulation and Signaling in Lung Adenocarcinoma. Cancer Res, 2018. 78(17): p. 4984–4996.

76. Zhang, S., et al., NUSAP1 is Upregulated by Estrogen to Promote Lung Adenocarcinoma Growth and Serves as a Therapeutic Target. Int J Biol Sci, 2024. 20(13): p. 5375–5395.

77. Ma, J., et al., Comprehensive multi-omics analysis identifies NUSAP1 as a potential prognostic and immunotherapeutic marker for lung adenocarcinoma. Int J Med Sci, 2025. 22(2): p. 328–340.

78. Xu, Z., et al., NUSAP1 knockdown inhibits cell growth and metastasis of non-small-cell lung cancer through regulating BTG2/PI3K/Akt signaling. J Cell Physiol, 2020. 235(4): p. 3886–3893.

79. Pilyugin, M., et al., BARD1 serum autoantibodies for the detection of lung cancer. PLoS One, 2017. 12(8): p. e0182356.

80. Zhang, Y.Q., et al., BARD1: an independent predictor of survival in non-small cell lung cancer. Int J Cancer, 2012. 131(1): p. 83–94.

81. Wang, Y., et al., USP4 function and multifaceted roles in cancer: a possible and potential therapeutic target. Cancer Cell Int, 2020. 20: p. 298.

82. Das, T., et al., USP15 and USP4 facilitate lung cancer cell proliferation by regulating the alternative splicing of SRSF1. Cell Death Discov, 2022. 8(1): p. 24.

83. Zhong, M., Q. Jiang, and R. Jin, USP4 expression independently predicts favorable survival in lung adenocarcinoma. IUBMB Life, 2018. 70(7): p. 670–677.

84. Wei, Y., et al., USP4 promotes proliferation and metastasis in human lung adenocarcinoma. Sci Rep, 2025. 15(1): p. 11096.

85. Yu, M., et al., Structural insight into ASH1L PHD finger recognizing methylated histone H3K4 and promoting cell growth in prostate cancer. Front Oncol, 2022. 12: p. 906807.

86. Xie, M., et al., The ASH1L-AS1-ASH1L axis controls NME1-mediated activation of the RAS signaling in gastric cancer. Oncogene, 2023. 42(46): p. 3435–3445.

87. Mäki-Nevala, S., et al., Driver Gene and Novel Mutations in Asbestos-Exposed Lung Adenocarcinoma and Malignant Mesothelioma Detected by Exome Sequencing. Lung, 2016. 194(1): p. 125–35.

88. Li, M., et al., Exploring the correlation between Tom1L1 and the efficacy of neoadjuvant chemotherapy for locally progressive mid-low rectal cancer. BMC Cancer, 2024. 24(1): p. 1413.

89. Wang, L., et al., TOM1L1 mediated the sort of tumor suppressive miR-378a-3p into exosomes and the excretion out of cells to promote ESCC progression. Cancer Gene Ther, 2025. 32(5): p. 507–520.

90. Chevalier, C., S. Roche, and C. Benistant, Vesicular trafficking regulators are new players in breast cancer progression: Role of TOM1L1 in ERBB2-dependent invasion. Mol Cell Oncol, 2016. 3(4): p. e1182241.

91. Chevalier, C., et al., TOM1L1 drives membrane delivery of MT1-MMP to promote ERBB2-induced breast cancer cell invasion. Nat Commun, 2016. 7: p. 10765.

92. Ding, S., et al., Identification of Pan-Cancer Biomarkers Based on the Gene Expression Profiles of Cancer Cell Lines. Front Cell Dev Biol, 2021. 9: p. 781285.

93. Sun, T., et al., Midazolam increases cisplatin-sensitivity in non-small cell lung cancer (NSCLC) via the miR-194-5p/HOOK3 axis. Cancer Cell Int, 2021. 21(1): p. 401.

94. Yang, K., et al., HOOK3 suppresses proliferation and metastasis in gastric cancer via the SP1/VEGFA axis. Cell Death Discov, 2024. 10(1): p. 33.

95. Melling, N., et al., High-Level HOOK3 Expression Is an Independent Predictor of Poor Prognosis Associated with Genomic Instability in Prostate Cancer. PLoS One, 2015. 10(7): p. e0134614.

96. Tan, E.H., et al., A multicentre phase II gene expression profiling study of putative relationships between tumour biomarkers and clinical response with erlotinib in non-small-cell lung cancer. Ann Oncol, 2010. 21(2): p. 217–222.

97. Zhang, J., et al., Has_circ_ASH1L acts as a sponge for miR-1254 to promote the malignant progression of cervical cancer by targeting CD36. Cancer Gene Ther, 2025. 32(2): p. 214–226.

98. Feng, Z.H., et al., EIF4A3-induced circular RNA PRKAR1B promotes osteosarcoma progression by miR- 361-3p-mediated induction of FZD4 expression. Cell Death Dis, 2021. 12(11): p. 1025.

99. Liu, G., et al., E2F3 promotes liver cancer progression under the regulation of circ-PRKAR1B. Mol Ther Nucleic Acids, 2021. 26: p. 104–113.

100. Cheng, D., et al., Downregulation of circ-RAPGEF5 inhibits colorectal cancer progression by reducing the expression of polypeptide N-acetylgalactosaminyltransferase 3 (GALNT3). Environ Toxicol, 2024. 39(8): p. 4249–4260.

101. Qiu, S., et al., Adult-onset CNS myelin sulfatide deficiency is sufficient to cause Alzheimer’s disease- like neuroinflammation and cognitive impairment. Mol Neurodegener, 2021. 16(1): p. 64.

102. Dobin, A., et al., STAR: ultrafast universal RNA-seq aligner. Bioinformatics, 2013. 29(1): p. 15–21.

103. Nielsen, A.F., et al., Best practice standards for circular RNA research. Nat Methods, 2022. 19(10): p. 1208–1220.

104. Vromman, M., J. Vandesompele, and P.J. Volders, Closing the circle: current state and perspectives of circular RNA databases. Brief Bioinform, 2021. 22(1): p. 288–297.

105. Chen, T., et al., The Genome Sequence Archive Family: Toward Explosive Data Growth and Diverse Data Types. Genomics Proteomics Bioinformatics, 2021.

106. Database Resources of the National Genomics Data Center, China National Center for Bioinformation in 2024. Nucleic Acids Res, 2024. 52(D1): p. D18–d32.

