## Supplemental methods for "Noncanonical Circular RNAs and Potential Functions"

### 1. Circular RNA Detection Using the CAT Pipeline

#### 1.1. Overview of the CAT algorithm

CircRNA of All Types (CAT) is a pipeline designed to comprehensively detect circular RNAs (circRNAs) from RNA-seq data without relying on genome annotation. This annotation-free approach enables the identification of both boundary and nonboundary circRNAs originating from genomic loci across the entire genome. The pipeline consists of three major steps: (1) initial mapping to the reference genome, (2) back-splicing junction (BSJ) detection by splitting mapping, (3) classification, expression quantification and filtering of selected circRNAs.

##### *1.1.1. Step 1: Initial mapping to reference genome*

RNA-seq reads are mapped to the reference genome using STAR v2.7.9a with the following parameters: --quantMode TranscriptomeSAM --outFilterMismatchNmax [kmis] --outReadsUnmapped Fastx --alignEndsType EndToEnd --outSAMunmapped Within. The mismatch tolerance parameter kmis is set based on read length, allowing one mismatch per 100 nucleotides. For reads of length 150 bp, kmis is set to 1. This conservative mismatch threshold is chosen to minimize false positives while still accommodating sequencing errors. The pipeline retains both mapped reads (for expression quantification) and unmapped reads (for circRNA detection) in separate files.

##### *1.1.2. Step 2: Candidate Back-splicing junction detection*

This step starts with the split mapping of unmapped reads to the genome. Short *anchor sequences* are first extracted from both ends of an unmapped read. Based on some empirical testing, the anchor length is set at one-fifth of the read length, with the maximum capped at 40 bp, to balance specificity (longer anchors map more uniquely) and sensitivity (shorter anchors are less affected by junction sequences). For example, for 150 bp reads, 30 bp anchors are extracted from both the 5' and 3' ends of every unmapped read. This is done by using the seqkit tool to extract the appropriate subsequences from every unmapped read.

The anchors are then mapped to the reference genome using Bowtie2 v2.4.4 with the parameters: -f -p [threads] -x [bt2_dna_ref] -k 1 -1 [5prime_anchors] -2 [3prime_anchors] -S [output_sam] --score-min C,-6,0. The -k 1 parameter is used to report only the best alignment for each anchor, reducing computational complexity while maintaining high confidence at the mapped location. The score threshold -- score-min C,0,0 is selected to disallow mismatch within the anchor.

The resulting SAM files are parsed to identify reads with both anchors mapped successfully to different genomic loci. Every genome-mapped anchor pair is then processed to find candidate back-splicing junctions, as follows:

1. Determine the genomic coordinates and strands of both anchors,
2. Verify that the orientation of the anchors is consistent with a circular structure (i.e., the 5' anchor mapped downstream of the 3' anchor on the same strand),
3. Extend the two anchors toward each other along the read sequence while maintaining alignment with their respective genomic loci,
4. Identify the precise BSJ point where the sequence transitioned from one genomic location to another, and

The anchor-extension process continued until either a BSJ point is detected or the number of mismatches exceeds the threshold of $\lfloor L/100 \rfloor$ for reads with L pb. This threshold is chosen to scale with read length, accommodating the higher probability of sequencing errors in longer reads while maintaining stringent quality control.

#### ***1.1.3. Step 3: Validation, expression quantification, filtering and classification***

##### ***Validation and quantification***

To validate candidate BSJs and quantify circRNA expression, a custom reference containing the sequences of candidate junctions detected so far was created by extracting genomic sequences spanning each candidate junction with the remap_ref function. Specifically, we extracted 150 bp (equivalent to the read length of the sample used to detect circRNA) upstream of the 3' junction site and 150 bp downstream of the 5' junction site, creating a 300 bp linear reference that represented the circularized junction region. This reference contains the unique back-splicing junctions that distinguish circRNAs from linear transcripts. This reference was indexed using Bowtie2-build for efficient alignment. All unmapped reads from Step 1 were then realigned to this junction reference using Bowtie2 with the parameters: -p [threads] --seed 42 -x <remap_idx_path> -1 <unmapped_mate1> -2 <unmapped_mate2> -S <remap_res_dir> --score-min C,[min_score],0 --rfg 6,6. The min_score parameter was dynamically set based on read length (default: -6 for 100 bp reads), allowing one mismatch per 100 nucleotides. The --seed 42 parameter ensured reproducible results, while --rfg 6,6 optimized the gap penalties for detecting BSJs.

The resulting alignments were then used to count the number of reads supporting each BSJ. Only BSJs with read counts exceeding a predefined threshold (default: 5 reads) were retained for further analysis. This threshold was deliberately set low to maximize sensitivity, particularly for low-abundance circRNAs, while still filtering out potential artifacts arising from sequencing errors.

##### ***Filtering***

The pipeline applied several filtering criteria to ensure high-quality circRNA predictions:

1. Retain only circRNAs with both sides of their junctions covered by sequencing reads. This criterion ensured that each reported circRNA had sufficient sequencing evidence supporting both sides of its BSJ, reducing false positives from mapping artifacts.
2. Remove duplicate BSJs due to multiple supporting reads with slightly different mapping patterns.
3. Filter out circRNAs exceeding a maximum estimated size threshold (default: 50,000 bp). This is based on the observation that most circRNAs are under 50 kb, and larger candidates are more likely to represent artifacts or genomic rearrangements rather than true circular transcripts.

##### ***Classification***

The candidate circRNAs are then classified based on their BSJs' genomic loci in reference to the genome annotation specified in the genome GTF annotation file. Note that the genome annotation is used here to classify circRNAs rather than detect them. This classifies the already-detected circRNAs into subtypes:

- Boundary circRNAs: BSJs coincide with annotated exon-intron boundaries,
- Complete nonboundary circRNAs: BSJs locate within annotated exonic or intronic regions,
- Half boundary circRNAs: One end of BSJs appears at an annotated exon-intron boundary, and
- Intergenic circRNAs: BSJs are located within intergenic regions

##### *1.1.4. Implementation details*

The CAT pipeline was implemented in Python 3.7+ with alliance on several bioinformatics tools and libraries, including STAR, Bowtie2, seqkit, pandas, and tqdm. The pipeline was designed to be memory-efficient and parallelizable, with configurable thread usage (default: 100 threads) to optimize performance on high-performance computing environments. This high number of default threads was chosen to maximize processing speed on modern multi-core servers but can be adjusted downward for systems with fewer available cores.

For paired-end data, the pipeline processed both read pairs simultaneously, while for single-end data, it adapted the workflow accordingly. The pipeline also incorporated strand-specific information when available, using the parameter strand_specific which could be set to 'F2R1' (forward strand corresponds to read 2, reverse to read 1), 'F1R2' (forward strand corresponds to read 1, reverse to read 2), or '' (for non-strand-specific data). The default 'F2R1' setting corresponds to the most common strand-specific library preparation protocols.

The entire workflow was designed with checkpoints to avoid redundant computation, checking for the existence and size of output files before executing each step. This design allowed for the efficient resumption of interrupted analyses and facilitated iterative parameter optimization.

### 2. Gene Feature Analysis

To investigate the genomic features associated with circRNA production, we performed comprehensive analyses of genes that produce different types of circRNAs. The analysis was conducted using Python 3.7+ with pandas and numpy libraries for data processing, and the gtfparse package for parsing genome annotation files.

We first categorized genes into three distinct groups based on their circRNA production patterns: genes producing (1) only canonical circRNAs, (2) only noncanonical circRNAs, and (3) both types (shared genes). This classification allowed us to examine whether specific gene features correlate with the propensity to generate different types of circRNAs.

For each gene group, we extracted and analyzed several key genomic features using the human genome annotation (Ensembl GRCh38.108). The primary features examined included gene length (measured from transcription start to end sites) and the number of exons. For genes with multiple annotated transcripts, we selected the first transcript listed in the genome annotation to maintain consistency across the analysis.

To ensure accurate representation of circRNA-gene relationships, we cross-referenced the gene information with our circRNA detection results. This allowed us to associate each circRNA with its host gene and corresponding genomic features. For genes hosting multiple circRNAs, we recorded the number of distinct circRNAs produced by each gene, providing insights into the circRNA production capacity across different gene types.

Additionally, we performed KEGG pathway enrichment analysis using the gseapy package to identify biological pathways that might be enriched in genes producing specific types of circRNAs. The enrichment analysis was conducted separately for each gene group, with results ranked by statistical significance (-log10 adjusted p-value). This functional analysis provided insights into the potential biological roles and regulatory networks associated with genes producing different types of circRNAs.

### 3. Analysis of Conservation Across Species

The analysis used data from lung tissues of three species: humans, macaques, and mice. For each circRNA, we extracted a 50-bp sequence centered on the BSJ (25 bp from each side of the junction). This region was selected because it contains the critical sequence elements that define circRNA structure and formation. The sequence extraction was performed using the reference sequence field from the CAT output, which provides the genomic context of each circRNA junction.

To assess sequence similarity between circRNAs from different species, we employed the Needleman-Wunsch global alignment algorithm implemented in BioPython's pairwise2 module. The alignment was configured with a match score of 1 and no penalties for mismatches or gaps (globalxx mode), focusing primarily on sequence identity. This approach was chosen to maximize sensitivity for detecting evolutionarily related circRNAs despite potential sequence divergence across species.

The similarity score for a circRNA pair in two species was calculated as the ratio of matching positions to the length of the longer sequence, resulting in a normalized similarity metric ranging from 0 to 1. Two circRNAs were considered potentially homologous if they shared at least 80% sequence identity in their junction regions. This threshold was established based on empirical testing and consideration of typical sequence conservation rates across the species examined.

We also conducted a threshold sensitivity analysis by calculating the number of conserved circRNA pairs at different similarity thresholds ranging from 0.8 to 1.0. This analysis provided insights into the distribution of conservation levels and helped establish appropriate thresholds for defining truly conserved circRNAs.

### 4. Amplicon Sequencing Analysis

To experimentally validate the presence of predicted circular RNAs, we developed a specialized Amplicon sequencing analysis pipeline. This analysis attempts to provide direct evidence for the existence of BSJs that are characteristic of circular RNA molecules. The analysis is implemented in Python using pandas, BioPython, and subprocess modules to interface with external bioinformatics tools.

The validation began with the preparation of a custom BSJ sequence reference for circRNAs to be validated, which is the same as the BSJ sequence reference in Step 3 of CAT (Section 1.1.3 above). This custom BSJ reference is critical for detecting reads spanning the back-fusion junctions. For the alignment Amplicon-seq reads to the reference, we employed Bowtie2 with optimized parameters for sensitive local alignment. The method was configured to use the following parameters: --very-sensitive-local to maximize sensitivity for detecting junction-spanning reads. The analysis allowed one mismatch per read to account for sequencing errors while maintaining specificity.

### 5. Ribosome Profiling Analysis

To investigate the translational potential of circular RNAs, we developed a specialized ribosome profiling analysis pipeline. This approach allowed us to identify ribosome-protected fragments (RPFs) spanning circRNA back-fusion junctions, providing direct evidence for circRNA translation.

The pipeline began with rigorous quality control of raw ribosome profiling data. Input FASTQ files were processed using Trim Galore with parameters specifically optimized for ribosome-protected fragments: quality threshold (q) of 20, minimum length of 20 bp, and maximum length of 35 bp. These parameters were selected based on the characteristic size distribution of ribosome-protected fragments, which typically range from 28-30 pb in length. The quality control step ensured that only high-confidence reads were retained for downstream analysis.

Following quality control, the pipeline used a sequential filtering strategy to remove reads originating from various non-coding RNA species. This multi-step approach was designed to progressively enrich RPFs derived from translated RNAs. The filtering process utilized Bowtie2 with the parameter --score-min C,0,0 to ensure stringent mapping with no mismatches allowed. This parameter choice minimized false-positive alignments while maintaining high specificity for the target sequences.

The first filtering step removed reads mapping to ribosomal RNA (rRNA) using a comprehensive rRNA reference database. This step was crucial as rRNA typically constitutes a significant portion of ribosome profiling libraries. The command included the parameter --un-gz rmrRNA.fq.gz to retain unmapped reads for subsequent filtering steps.

The second filtering step targeted transfer RNA (tRNA) sequences, another major contaminant in ribosome profiling data. Reads that did not map to either rRNA or tRNA were then subjected to additional filtering against genomic DNA and RNA references to remove any remaining reads derived from linear transcripts.

After the sequential filtering process, the remaining reads represented potential RPFs from noncanonical RNA sources, including circular RNAs. These filtered reads were then mapped to a custom reference database of circRNA sequences. The reference database was created by extracting sequences spanning the BSJs of predicted circRNAs, with 150 bp long (Equivalent to the read length of the sample used to detect from each side of the junction points.

For the circRNA mapping step, we employed Bowtie2 with parameters optimized for detecting junction-spanning reads. The pipeline allowed one mismatch per read.

The results of the ribosome profiling analysis were compiled into detailed reports containing information on each translated circRNA, including its genomic coordinates, supporting read counts, and circRNA classification.

#### 6. Internal Ribosome Entry Site Analysis

To investigate the potential for cap-independent translation of circular RNAs into proteins or peptides, we analyzed Internal Ribosome Entry Site (IRES) elements within circRNA sequences.

The IRES detection pipeline used the circRNA BSJ reference sequences of the predicted circRNAs (see Section 1.1.3). For each circRNA, we utilized the full circular sequence reconstructed during the circRNA detection process, which included the genomic sequence spanning the BSJ. This ensured that potential IRES elements located near or spanning the junction would be properly identified.

Sequence similarity searches were performed using BLASTN (Basic Local Alignment Search Tool for nucleotides) to compare circRNA sequences against a curated database of experimentally validated IRES elements. The BLASTN parameters were optimized for IRES detection: -perc_identity 80 to require a minimum sequence identity of 80%, and -word_size 7 to increase sensitivity for shorter matches while maintaining specificity. These parameters were selected based on the characteristic features of functional IRES elements, which typically require conservation of specific structural motifs rather than perfect sequence identity.

Following the BLASTN search, we implemented a filtering step to identify high-confidence IRES elements within circRNAs. The filtering criteria required alignments to have Percent identity ⩾80% to ensure sufficient sequence similarity with known IRES elements, and Alignment length ⩾30 nucleotides to cover the minimal functional IRES region.

#### 7. Single-cell RNA-seq Analysis

We processed and analyzed a total RNA sample from lung cancer cells (A549 and HCC827) and leukemia cells (K562). Raw reads were first aligned and quantified using SeekSoulTools v1.2.2 and CellRanger v8.0.1, respectively, using the reference genome GRCh38. Each scRNA-seq library was then preprocessed to obtain data from high-quality cells. Specifically, for each library, the cell screening criteria included the number of expressed genes greater than 200 and less than 8500, the unique molecular identifier (UMI) count greater than 500 and less than top 0.5% total UMI number of each sample, the mitochondrial genes counted as less than 10% of the total UMI counts, and filtering genes that were expressed in less than three cells. Scrublet was used to estimate doublet probabilities and filter cells with scores greater than 0.3. To apply strict doublet filtering, the Scrublet workflow was also implemented.

Following preprocessing, the gene expression matrix of high-quality cells was obtained and used for clustering with the Scanpy v1.10.0 software. The clustering steps included (1) log-normalization with function of sc.pp.normalize_total (target_sum=1e4) followed by sc.pp.log1p(), (2) highly variable gene selection by function sc.pp.highly_variable_genes() with the flavor of seurat_v3 and choosing the top 3000 genes, (3) computing principal components by function sc.pp.pca() with first 20 principle components, (4) construction of a k nearest-neighbor (KNN) graph using function sc.pp.neighbors with 30 neighbors and the Euclidean distance, and (5) Leiden clustering using function sc.tl.leiden() with the resolution of 1. In addition, for visualization, uniform manifold approximation and projection (UMAP) for nonlinear dimensionality reduction was computed via the function sc.tl.umap(). These clusters were annotated using known cell-type marker genes, and noisy clusters and doublet clusters were identified and removed based on the low co-expression of these genes. For the cell line sample, marker genes included those for HCC827 (EGFR, EPCAM, NFKBIZ); A549 (ALDH1A1, AKR1C3, SERPINE1), and K562 (HBG1, HBG2, ANK1).

**Software Availability**

The complete analysis pipeline and associated scripts are available at [https://github.com/GenomicMedicine/CAT].
